# Neural correlates of modality-specific and modality-invariant object recognition in the perirhinal cortex

**DOI:** 10.1101/2023.11.20.567750

**Authors:** Heung-Yeol Lim, Inah Lee

## Abstract

The perirhinal cortex (PER) supports multimodal object recognition, but how multimodal information of objects is integrated within the PER remains unknown. Here, we recorded single units within the PER while rats performed a PER-dependent multimodal object-recognition task. In this task, audiovisual cues were simultaneously (multimodally) or separately (unimodally) presented. We identified two types of object-selective neurons in the PER: crossmodal cells, showing constant firing patterns for an object irrespective of its modality, and unimodal cells, showing a preference for a specific modality. Unimodal cells further dissociated unimodal and multimodal versions of the object by modulating their firing rates according to the modality condition. A population-decoding analysis confirmed that the PER could perform both modality-invariant and modality-specific object decoding – the former for recognizing an object as the same in various conditions and the latter for remembering modality-specific experiences of the same object.

## Introduction

Our brains can effortlessly integrate information from different sensory modalities to form a unified representation of the world ^1,2^. This natural ability is also evident during object recognition, as one can quickly identify one’s cat by visually perceiving its appearance or hearing its distinctive meow. The ability to recognize objects crossmodally has been reported not only in humans, but also in nonhuman primates ^3,4^, rodents ^5–7^, dolphins ^8^, and even insects ^9^. However, most studies on object recognition have neglected the multisensory nature of this process. Object recognition has been studied primarily using unimodal stimuli, such as visual stimuli ^10–12^, or uncontrolled multimodal stimuli, such as 3D “junk” objects ^13,14^, without a specific goal of investigating multimodal processing. This tendency is also evident in studies of the perirhinal cortex (PER), a region well known to play a critical role in object recognition ^15–20^.

Findings from several studies have implied that the PER is engaged in “multimodal” object recognition. Anatomically, it has been shown that the PER receives inputs from areas that process diverse sensory modalities, including those from visual, auditory, olfactory, and somatosensory cortices ^21–23^. In rodents in particular, these areas are known to send monosynaptic inputs to the PER ^22^. Experimental results further support the involvement of the PER in multimodal object recognition. In human functional magnetic resonance imaging (fMRI) studies in which subjects were presented visual-auditory or visual-tactile stimuli that were either from the same (congruent) or different (incongruent) objects, activity within the PER was found to be greater when the two stimuli were congruent ^24,25^. The necessity of the PER for multimodal object recognition has also been tested using crossmodal versions of a delayed nonmatch-to-sample task in nonhuman primates ^4^ and a spontaneous object-recognition task in rodents ^5–7^. In these tasks, in which animals sampled an object using one sensory modality (e.g., tactile), and then were tested for retrieval of object information using an unused sensory modality (e.g., visual), lesioning or inactivating the PER resulted in performance deficits. These results indicate the involvement of the PER in multimodal object recognition, but the mechanisms underlying these functions remain largely unknown.

We hypothesized that the PER may support multisensory object recognition by integrating multimodal inputs from an object to form a unified representation of the object. Considering the associative nature of the PER ^26–29^, the region can be expected to integrate information from multiple sensations, rather than processing it separately. Indeed, it has been shown that PER neurons do not represent individual sensory attributes separately in rats performing behavioral tasks using multimodal stimuli ^30,31^. However, these studies have only reported neural correlates of behavioral responses or rewards associated with objects, rather than actual information about the objects themselves. Accordingly, in the current study, we investigated how multimodal information is integrated to create a unified representation of an object while minimizing the influence of other task-related variables, such as behavioral response or reward outcome.

To test the abovementioned hypothesis, we developed a multimodal object-recognition task for rats employing visual and auditory cues. By requiring a nose-poke during object cue sampling, our task allowed presentation of different task phases while observing their neural firing correlates in a temporally controlled manner. Our findings suggest that rats can recognize a familiar object (originally learned multimodally) almost immediately when cued by a unimodal sensory attribute alone (e.g., visual or auditory) without additional learning. However, inactivating the PER resulted in performance deficits in both multimodal and unimodal recognition conditions. Physiologically, we discovered that most PER neurons exhibited a constant selectivity pattern for an object regardless of its sensory modality. However, a significant proportion of neurons also showed a preference for a specific sensory modality condition during object information processing. A population-decoding analysis revealed that these subpopulations of neurons enabled both modality-specific and modality-invariant recognition of objects.

## Results

### The PER is required for multimodal object recognition

To test multimodal object recognition while controlling the sampling of the object’s unimodal (i.e., visual and auditory) attributes, we developed a behavioral paradigm for rats that would enable stable, simultaneous sampling of multimodal cues (**Fig. 1A**). In the sample phase of this protocol, rats triggered the onset of cues by nose-poking a center hole and were required to maintain their nose-poke for at least 400 ms. If a rat failed to maintain its nose-poke for 400 ms, the trial was stopped and the rat was allowed to retry the nose-poke after a 4-s interval (**Fig. S1**). After a successful (>400 ms) nose-poke, the cues disappeared and doors covering left and right choice ports were opened simultaneously. In the response phase, rats were required to choose either the left or right port based on the sampled cue. Rats completed their choice responses within 600 ms in most trials (**Fig. S2**). A food reward was provided only after a correct choice response was made (reward phase), followed by 2-s inter-trial interval.

**Fig. 1.**
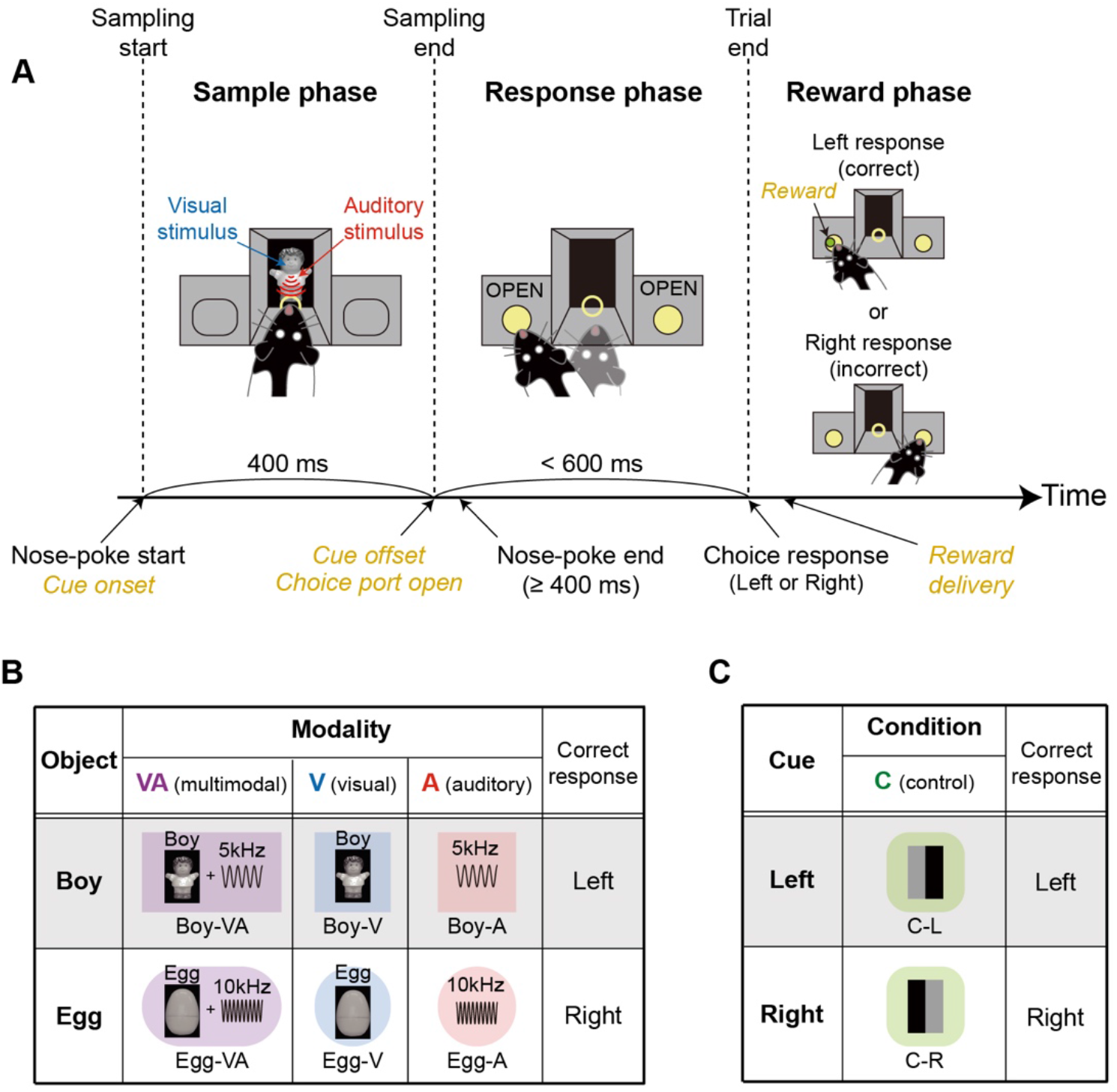
Multimodal object-recognition task. (**A**) Illustration of the apparatus and the trial structure of the multimodal object-recognition task. Rats sampled visual and auditory cues simultaneously or separately for 400 ms (sample phase) and then made a choice response based on the identity of the cue (response phase). A correct choice response resulted in a food reward (reward phase). (**B**) Object conditions used in the multimodal object recognition task. Two different objects (*Boy* and *Egg*) were presented in three different modality conditions: multimodal (VA), visual (V), and auditory (A). The correct choice response was determined by the identity of the object. (**C**) Two simple visual cues were introduced as control (C) stimuli. Each control stimulus was also associated with either the left (C-L) or right (C-R) choice response (i.e., the same responses required by object conditions).

To test the rat’s ability to recognize objects with multiple sensory modalities, we presented two different multimodal objects, *Boy* and *Egg*, consisting of different combinations of visual (images of a boy-shaped and an egg-shaped toy) and auditory (5 and 10 kHz sine-wave tones) attributes during the sample phase (**Fig. 1B**). Objects were tested under three modality conditions: multimodal, visual, and auditory. In the multimodal condition, visual and auditory cues associated with an object were presented simultaneously during the sample phase. In unimodal – visual or auditory – conditions, only the object’s visual or auditory information was presented as a cueing stimulus. If the rat responded correctly to the object’s identity regardless of the modality condition, it was rewarded with a piece of cereal. The combination of audiovisual cues and stimulus-response contingency were counterbalanced across rats. In control conditions, rats learned to dissociate two simple visual stimuli composed of black and gray bars (**Fig. 1C**). In these conditions, the required left and right choice responses were the same as those in object conditions. In sum, eight stimulus conditions were used in this task: six object conditions (two objects × three modality conditions) and two control conditions.

To test whether rats are able to retrieve multimodal objects when cued by a unimodal stimulus under conditions in which the PER is inactivated, we conducted a drug-inactivation experiment (n = 6). After training in multimodal and control conditions, rats were sequentially tested under multimodal, visual, auditory, and control conditions in separate sessions (**Fig. 2A**). The order of visual and auditory sessions was counterbalanced across rats. For each condition, we first established baseline performance by injecting vehicle control (phosphate-buffered saline [PBS]) into the PER; we then tested performance in rats with an inactivated PER, achieved by injecting muscimol (MUS) bilaterally into the PER. Importantly, the sessions with PBS injections, either visual (V1) or auditory (A1) (**Fig. 2A**), marked the first instances where rats were required to recognize objects, originally learned multimodally, solely based on their unimodal sensory attributes. In a unimodal object recognition session, objects were presented multimodally (visual and auditory) for the first 20 trials, and then subsequently presented in a unimodal (visual or auditory) fashion.

**Fig. 2.**
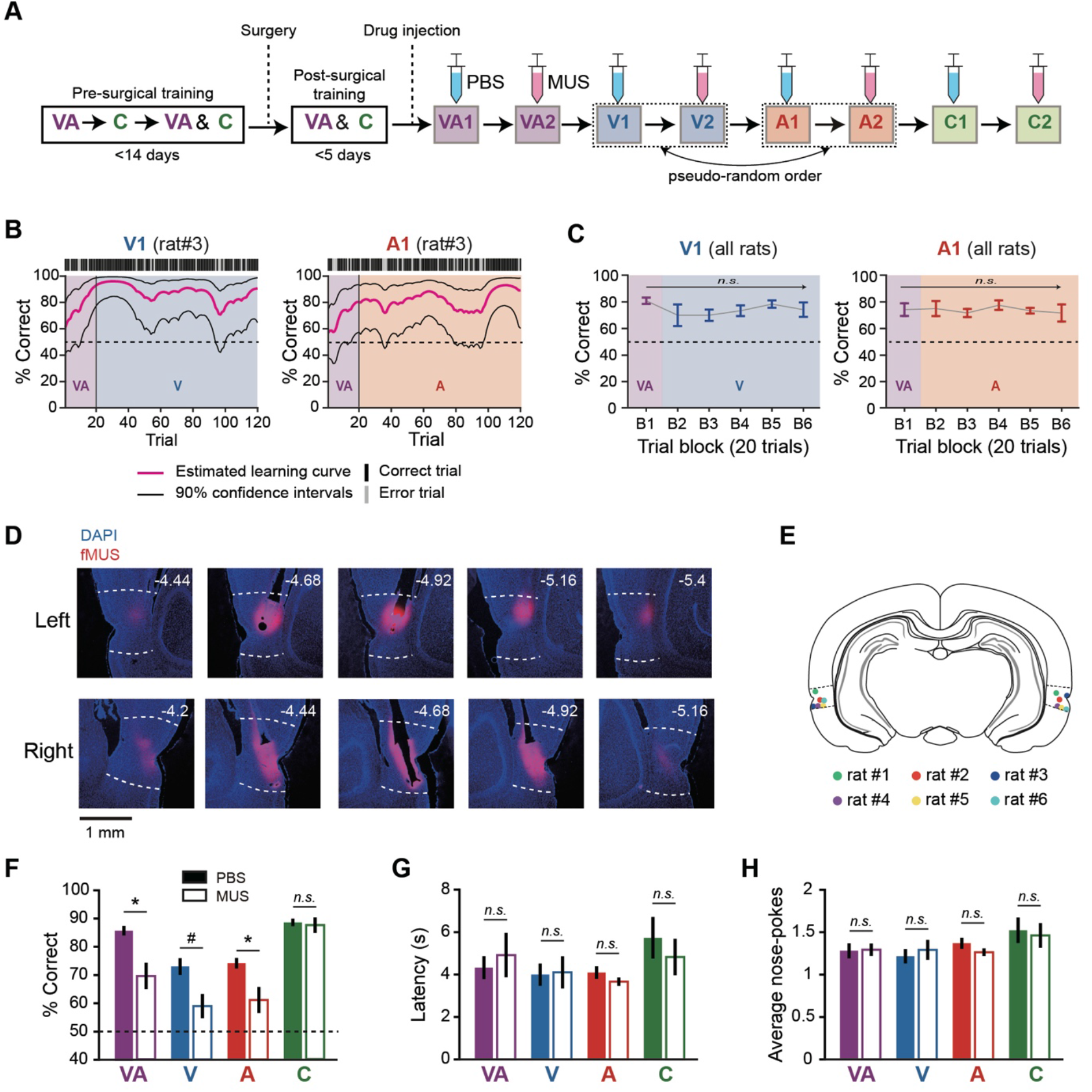
Necessity of the PER for multimodal object recognition. (**A**) Illustration of behavioral training and testing schedules for the PER-inactivation experiment. Note that animals were subjected to either the visual or auditory condition for the first time in PBS-injected visual (V1) or auditory (A1) sessions. (**B**) Estimated learning in V1 (left) and A1 (right) sessions of an example rat. In trial 21, where visual or auditory conditions were first introduced, the rat quickly adapted without additional learning. (**C**) On average, correctness did not significantly change across trials within V1 (left) or A1 (right) session, indicating that rats could perform unimodal retrieval without additional learning. Each trial block consisted of 20 trials. (**D**) Histological verification of injection sites in the PER. White dotted lines indicate the border of the PER. The numbers on each section indicate the distance from bregma. (**E**) Summary of cannula-tip locations in all rats. (**F**) Behavioral performance in each condition was compared between PBS and MUS sessions. Performance was significantly impaired in all object conditions (VA, V, and A) by inactivation of the PER, but remained intact in the control (C) condition. (**G**) The latency median did not change significantly after inactivating the PER. (**H**) The average number of nose-poke attempts did not change significantly after inactivating the PER. Data are presented as means ± SEM (n = 6; \**p* < 0.05, ^#^*p* = 0.062; n.s., not significant).

Performance dynamics of PBS-injected rats in visual and auditory sessions were displayed as learning curves, estimated from a given session (**Fig. 2B**). Upon first encountering the visual or auditory condition (Trial 21), rats showed no significant drop in performance and their performance remained stable until the end of the session. A statistical analysis of results for all PBS-injected rats revealed no significant increase or decrease in performance across trial blocks (20 trials) in either visual (F_(5,25)_ = 0.95, *p* = 0.47) or auditory (F_(5,25)_ = 0.22, *p* = 0.95; one-way repeated measures ANOVA) sessions (**Fig. 2C**). These results indicate that rats easily recognized an object originally learned multimodally using one of its unimodal attributes, and this crossmodal recognition process required minimal training.

To verify the necessity of the PER in the task, we examined the effect of MUS injection on task performance. Histological results confirmed that MUS was successfully bilaterally injected into the PER (**Fig. 2D** and **2E**). The average performance of rats (n = 6) in PBS sessions was significantly higher than predicted by chance (50%) in all conditions – multimodal (t_(5)_ = 21.2 *p* < 0.0001); visual (t_(5)_ = 7.8, *p* = 0.0005); auditory (t_(5)_ = 13.1, *p* < 0.0001); and control (t_(5)_ = 29.3, *p* < 0.0001) – as determined by one-sample t-test. Inactivating the PER with MUS significantly decreased performance (F_(1,5)_ = 165.4, *p* = 0.0006, two-way repeated measures ANOVA) (**Fig. 2F**). A post hoc analysis revealed performance deficits in multimodal (t_(5)_ = 3.72, *p* = 0.028), visual (t_(5)_ = 2.39, *p* = 0.062), and auditory (t_(5)_ = 3.45, *p* = 0.027) conditions (paired t-test with Holm-Bonferroni correction), but not in the control condition (t_(5)_ = 0.37, *p* = 0.36, paired t-test). Trial latency (i.e., from trial onset to end of choice) was not significantly affected by MUS injection (F_(1,5)_ = 0.13, *p* = 0.73; two-way repeated measures ANOVA) (**Fig. 2G**). Nose-poking behavior was not affected by PER inactivation, as the average number of nose-poke attempts was not significantly different between PBS and MUS sessions (F_(1,5)_ = 0.92, *p* = 0.38, two-way repeated measures ANOVA) (**Fig. 2H**). Collectively, these results demonstrate that the PER is necessary for object recognition in all modality conditions and that the decrease in performance is not attributable to a generic deficit.

### Object-selective neural activity in the PER is characterized by its transient and sequential firing patterns

Inactivation of the PER resulted in performance deficits whenever object recognition was required regardless of the modality condition. To further understand the functions of the PER in multimodal object recognition, we searched for neural correlates of multimodal object recognition by recording single-unit spiking activity in the PER using tetrodes (**Fig. 3A**). Based on their basic firing properties, most neurons could be classified into regular-spiking neurons (68%, 234 of 348), with bursting (24%, 82 of 348) and unclassified (9%, 32 of 348) neurons also being observed (**Fig. 3B**), as previously reported ^16,32^.

**Fig. 3.**
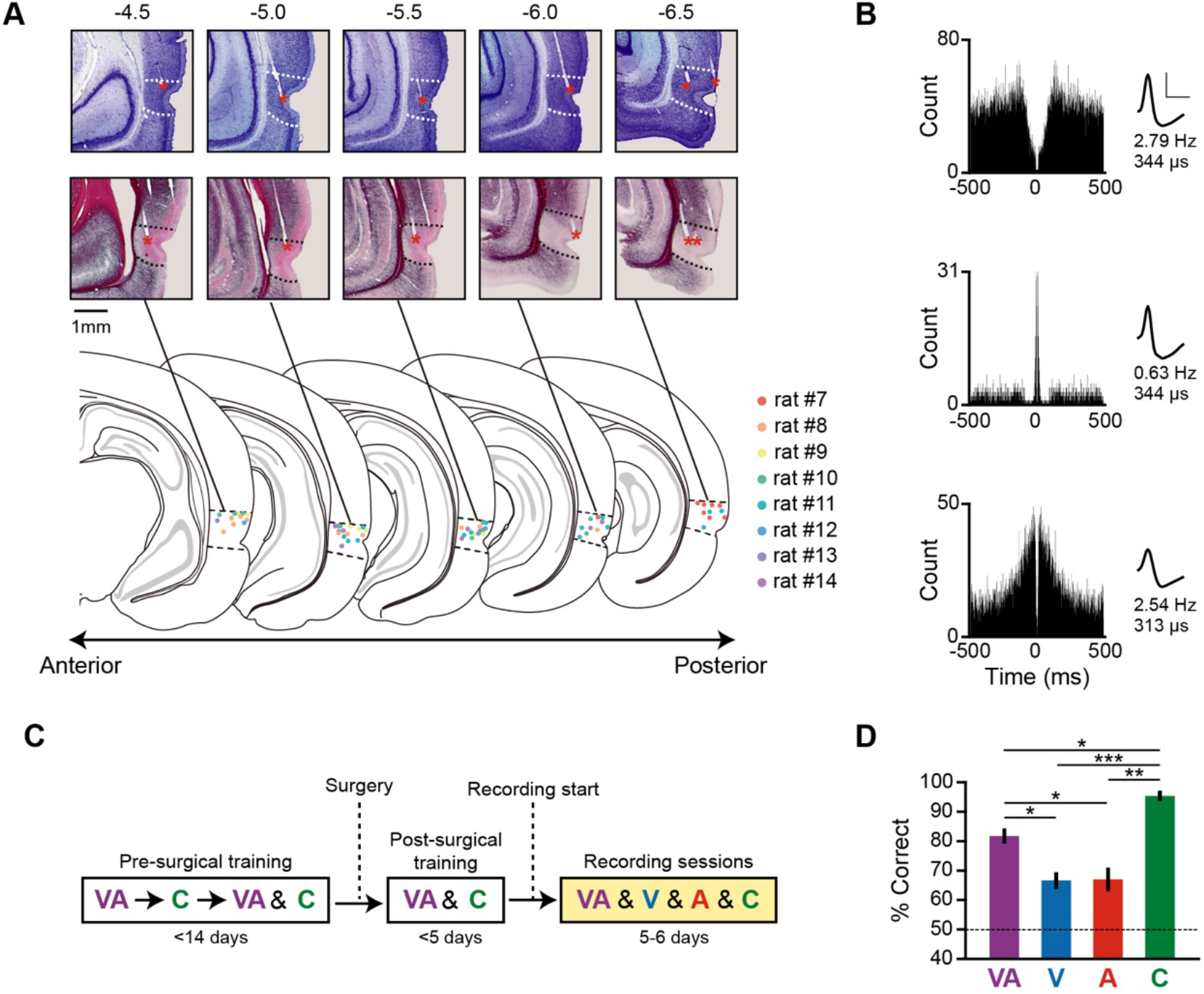
Single-neuron recordings during multimodal object recognition. (**A**) Histological verification of tetrode locations in the PER by Nissl (top) and myelin (middle) staining of sections across the anteroposterior axis. The estimated tetrode tip locations in all rats are summarized on the atlas (bottom). Dotted lines demarcate the borders of the PER. Tetrode tip locations are marked with red asterisks. The numbers above each section indicate the distance from bregma (mm). (**B**) Examples of single neurons classified according to their basic firing properties. Based on the autocorrelograms (left), cells were categorized as regular-spiking (top), bursting (middle), or unclassified (bottom). Scale bars in each spike waveform (right) indicate amplitude (vertical, 100 μV) and width (horizontal, 500 μs). The numbers below the waveform show the mean firing rate and spike width of each neuron. (**C**) Illustration of training and recording schedules for electrophysiological experiments. In the recording sessions, all stimulus conditions (VA, V, A, C) were pseudo-randomly presented within a session. Rats experienced visual or auditory conditions only in the recording sessions. (**D**) Behavioral performance in the first recording session. Although rats performed better in pre-trained multimodal and control conditions, they still showed better than chance-level performance in visual and auditory conditions. Data are presented as means ± SEM (n = 8; \**p* < 0.05, \**p* < 0.05, \*\**p* < 0.01, \*\*\**p* < 0.001; n.s., not significant.)

Before obtaining single-unit recordings, rats were first trained in multimodal and control conditions; unimodal (visual or auditory) recognition conditions were introduced upon initiation of recordings (**Fig. 3C**). All testing conditions (multimodal, visual, auditory, and control) were presented pseudo-randomly within a recording session. We confirmed that rats (n = 8) were able to successfully recognize objects in all conditions in their first recording session – multimodal (t_(7)_ = 12.36, *p* < 0.0001); visual (t_(7)_ = 5.88, *p* = 0.0006); auditory (t_(7)_ = 4.26, *p* = 0.0037); and control (t_(7)_ = 25.9, *p* < 0.0001) – as determined using one-sample t-test (chance level, 50%) (**Fig. 3D**). Significant differences in performance were also noted among conditions (F_(3,21)_ = 22.87, *p* < 0.0001, one-way repeated measures ANOVA), with rats performing significantly better in the multimodal condition than in either the visual (t_(7)_ = 3.43, *p* = 0.022) or auditory (t_(7)_ = 4.22, *p* = 0.016; paired t-test with Holm-Bonferroni correction) condition. Performance in the control condition was significantly higher than that in all other conditions (control vs. multimodal, t_(7)_ = 3.92, *p* = 0.017; control vs. visual, t_(7)_ = 15.47, *p* < 0.0001; control vs. auditory, t_(7)_ = 6.19, *p* = 0.0023; paired t-test with Holm-Bonferroni correction). Similar behavioral results were observed in all recording sessions (**Fig. S2**).

We next sought to describe object selectivity of PER cells by determining how these neurons respond to different object identities regardless of sensory modality. To this end, we grouped all correct trials into different object and modality conditions and then calculated the firing rates associated with each condition during the task epoch, measured from the start of the sample phase to the end of the response phase (900-ms duration) (**Fig. 4A**). Overall firing patterns were obtained by averaging firing rates in different modality conditions for each object, *Boy* and *Egg* (**Fig. 4A** and **4B**, black lines). For each neuron, we defined an object-selective epoch as the period in which the firing rate for either object was significantly different from that of the other object in more than five consecutive time bins (10 ms/bin) (**Fig. 4B**, example neurons #1–6). Since the object-selective epoch defined here could be attributable to the choice response and not necessarily to the identity of the object, we further excluded response-selective cells identified under control condition and considered the remaining neurons to be object-selective cells (hereafter, object cells) (**Fig. S4**). Selectivity was not maintained throughout sample and response phases; thus, individual object cells were characterized by their transient firing patterns. Moreover, the time bin at which the firing rate difference between objects was maximal (i.e., peak selectivity time) occurred at various time points during the task epoch (**Fig. 4B**).

**Fig. 4.**
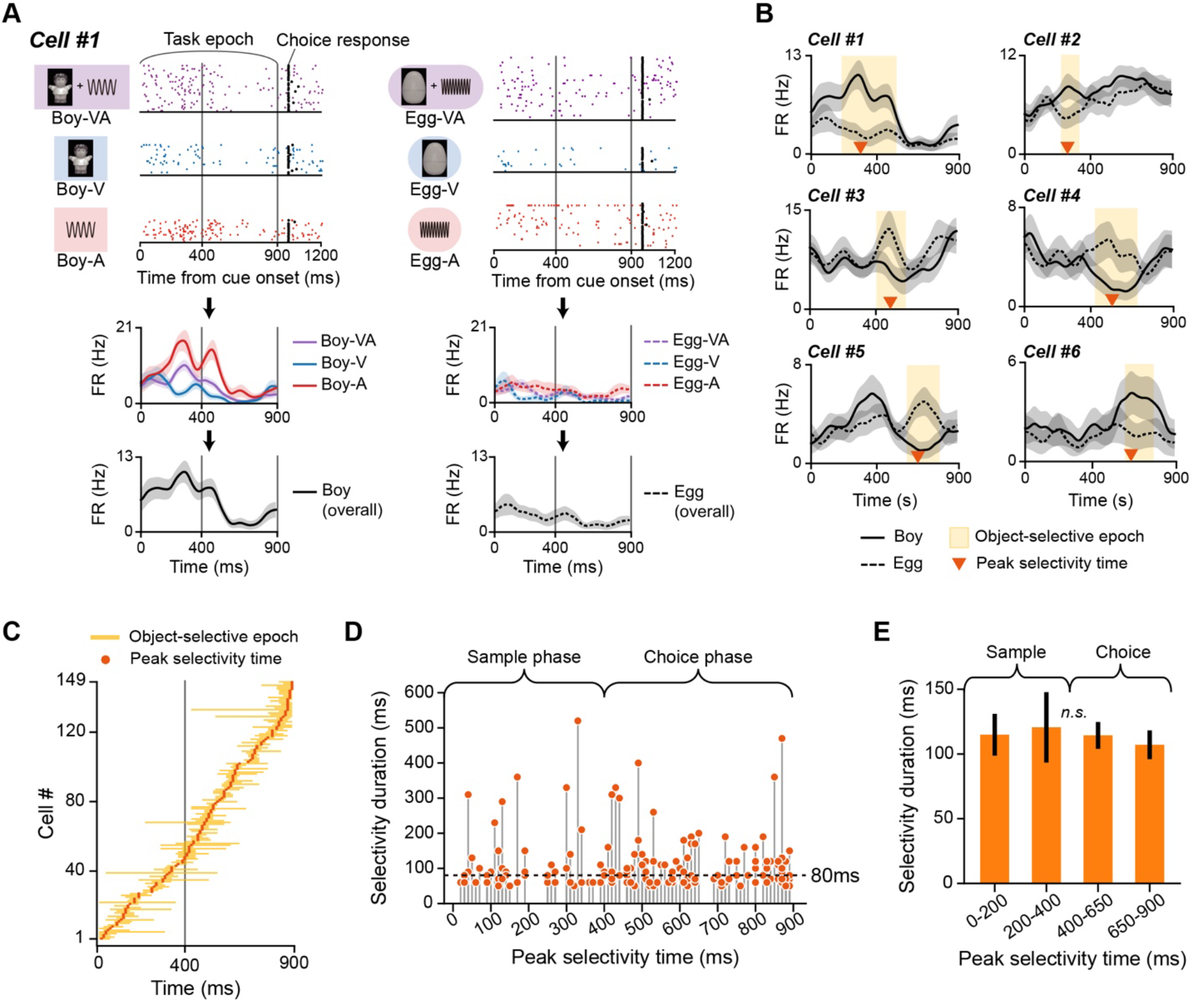
Object-selective firing patterns in the PER. (**A**) Raster plots (top) and spike density functions (bottom) of an example neuron for *Boy* (left) and *Egg* (right) object conditions. Overall firing rates for each object (black line) were obtained by averaging firing rates in different modality conditions (VA, V, and A). This sample neuron showed increased firing rates for the *Boy*, but not the *Egg* object (i.e., *Boy*-preferring neuron). Note that the interval from 0 to 900 ms after the cue onset, designated the task epoch, was the analysis target. (**B**) Example object cells in the PER showing selective firing patterns for an object over the object-selective epoch, indicated in yellow. Orange arrowheads indicate the peak selectivity time (i.e., time when selectivity was maximal). (**C**) Population object selectivity of all object cells and their peak selectivity times. The selective epoch of each object cell was marked and then aligned according to their peak selectivity time. The vertical gray line indicates the temporal boundary of sample and response phases. (**D**) Peak selectivity time and duration of the selective epoch. Each dot indicates an individual object cell. Dotted line denotes the median selectivity duration (80 ms). (**H**) Comparison of selectivity durations between cells whose peak selectivity times appeared in different time ranges. No significant difference was found. Data are presented as means ± SEM (n.s., not significant).

To visualize the characteristics of object cells at the population level, we constructed a population object-selectivity plot (**Fig. 4C**), in which object-selective epochs of individual object cells were marked and then aligned by their peak selectivity time. Interestingly, we observed a sequentially ordered firing of object-selective cells such that the population of object cells tiled the task epoch (from the sample phase to the response phase) with their object selectivity. We further investigated the possibility that object selectivity might be stronger in certain time bins, even when this sequential pattern was present. For this, we used the duration of selectivity as a measure of the magnitude of object selectivity and examined the relationship between the selectivity duration and peak selectivity time (**Fig. 4D**). The median selectivity duration was 80 ms, confirming the transient nature of object-selective firing in the PER. We found no evidence that cells with greater selectivity were more active in certain time bins. Selectivity durations were not significantly different upon grouping cells into four temporal intervals based on their peak selectivity time (F_(3,145)_ = 0.14, *p* = 0.96; one-way ANOVA) (**Fig. 4E**). Taken together, these observations indicate that object cells in the PER are characterized by their transient and sequential activity patterns, which tiled the entire task epoch. Notably, these characteristics were present regardless of whether the rats were sampling the cues (sample phase) or choosing a behavioral response in the absence of cues (response phase).

### Both visual and auditory information processings occur during object-selectivity firing in the PER

If PER neurons solely focus on the identity of an object and its associated behavioral response, object-selective patterns should remain constant irrespective of the modality condition. Conversely, it could be argued that distinguishing between events associated with experiencing an object based on its distinct modality information is crucial for episodic memory. To determine whether PER object cells can encode a particular sensory modality, we applied multiple linear regression to firing rates during the object-selective epoch (see Methods for details). In this regression model, *β_1_* and *β_2_* are regression coefficients that represent the visual and auditory responsiveness, respectively, of the preferred object (i.e., the object condition with higher firing rates). Visual and auditory information-processing neurons within the PER were identified based on the relationship between *β_1_* and *β_2_* (**Fig. 5A**). An example of an object cell that predominantly fired for the visual attribute of *Boy* is cell #7 (**Fig. 5A-ii**), which had higher firing rates in multimodal and visual conditions compared with the auditory condition. This pattern is reflected in higher *β_1_* versus *β_2_* values (**Fig. 5A-iii**). Cell #8, on the other hand, was responsive to the auditory attribute of *Boy*, as its firing rates in the multimodal and visual condition were higher compared with those in the visual condition (**Fig. 5A-ii**); it also had higher *β_2_* than *β_1_* values (**Fig. 5A-iii**). A crossmodal cell type, distinct from the unimodal cell type described above that exhibited no significant preference for a particular sensory modality, was also observed (**Fig. 5B**). An example of a crossmodal cell is cell #9, which exhibited almost equal firing in response to both sensory modalities of its preferred object (*Boy*) (**Fig. 5B-ii**); its *β_1_* and *β_2_* values were also similar (**Fig. 5B-iii**).

**Fig. 5.**
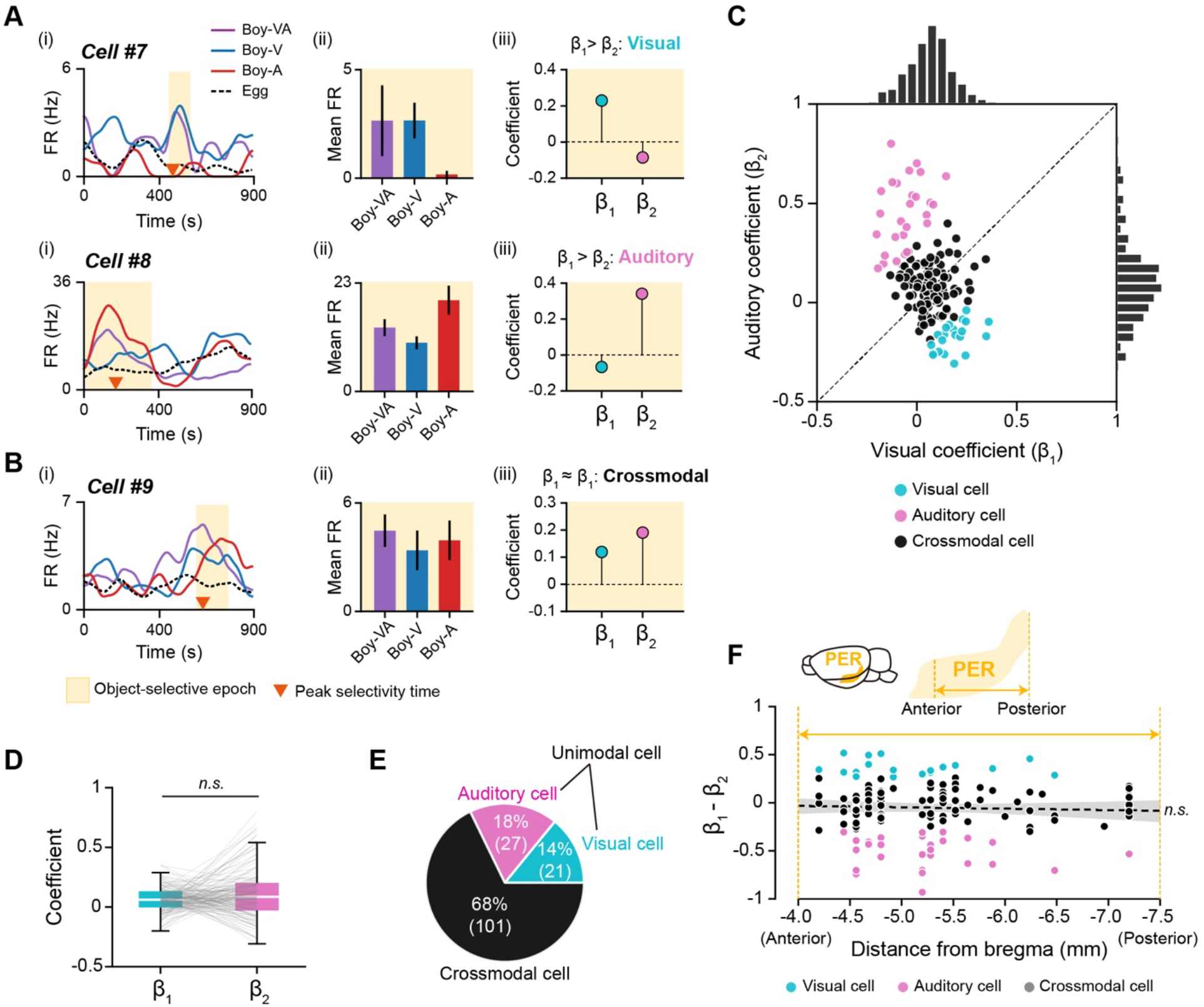
Unimodal and crossmodal response patterns of object cells in the PER. (**A**) Examples of unimodal cells that were responsive to either the visual or auditory attribute of an object during the selective epoch. Spike density functions (i) and mean firing rates within the object-selective epoch (ii). Multiple linear regression was applied to firing rates within the object-selective epoch to obtain β_1_ and β_2_ – regression coefficients reflecting the magnitude of visual and auditory responses, respectively (iii). Cell #7 mainly responded to the visual attribute of *Boy* (β_1_ > β_2_), whereas cell #9 was responsive to the auditory attribute of *Boy* (β_1_ < β_2_). (**B**) Spike density functions (i), mean firing rates (ii), and regression coefficients (iii) of a crossmodal cell. The cell showed no specific bias for visual or auditory information processing, as indicated by similar β_1_ and β_2_ values. (**C**) Scatter plot and histograms of visual (β_1_) and auditory (β_2_) coefficients in all object cells. Neurons were classified as either visual (cyan) or auditory (pink) cells if the difference between visual and auditory coefficient was significant. Others were classified as crossmodal cells (gray). (**D**) Visual and auditory coefficients of all object-selective cells were not significantly different. Each line indicates an individual object cell. (**E**) Proportions of visual, auditory, and crossmodal neurons within the object cell category. Visual and auditory cells were grouped as a unimodal cell type. The numbers in parentheses denote the number of neurons. (**F**) Anatomical locations of object cells along the anteroposterior axis of the PER and their unimodal (or crossmodal) response patterns. Differences between β_1_ and β_2_ did not exhibit a significant linear relationship with anatomical locations of the cells. The dotted black line indicates the linear regression line, and the shaded area is the 95% confidence interval (n.s., not significant).

To illustrate the patterns of modality correlates, we created a scatter plot of *β_1_* and *β_2_* values for all object cells (**Fig. 5C**). We then verified that the PER system did not preferentially process one of the sensory modalities by first comparing *β_1_* and *β_2_* for all object cells (**Fig. 5D**). This analysis showed no significant difference between *β_1_* and *β_2_* (W = 4794, *p* = 0.13; Wilcoxon signed-rank test), indicating that the PER did not have a significant bias toward a specific sensory modality. We then classified neurons based on the difference between their *β_1_* and *β_2_* values such that neurons whose *β_1_* values were significantly higher than their *β_2_* values were classified as visual cells, whereas those with significantly higher *β_2_* than *β_1_* values were classified as auditory cells. Other object cells were classified as crossmodal cells. Although the majority of object cells were categorized as crossmodal (68%), both auditory cells (18%) and visual cells (14%) were identified (**Fig. 5E**). The small difference in the proportion of visual and auditory cell categories was determined to be insignificant (χ^2^ = 0.89, *p* = 0.34; chi-square test). Detailed comparisons of selectivity patterns revealed that auditory cells exhibited stronger selectivity in the sample phase and their selective period was longer than that of visual cells (**Fig. S5**). These findings suggest that modality information processing within the PER is heterogeneous, potentially enabling the retrieval of both object identity and its associated modality information.

Since the PER receives direct inputs from visual and auditory cortices ^22,23^, it is possible that the activity of visual and auditory cells in the PER is driven solely by inputs from the sensory cortices. If so, the posterior PER, where visual inputs are relatively dominant, might have more visual cells, whereas the anterior PER, which receives more auditory inputs, might possess more auditory cells. To test this hypothesis, we examined the relationship between the anatomical locations of cells along the anteroposterior axis of the PER and differences between visual (*β_1_*) and auditory (*β_2_*) coefficients (**Fig. 5F**). We found no evidence for regional bias in coefficients in the posterior PER that would indicate the dominance of visual processing over auditory processing. Instead, visual and auditory cell types were evenly distributed along the anteroposterior axis of the PER. These results suggest that the activities of visual and auditory cells in the PER do not solely rely on inputs from visual and auditory cortices, respectively.

### Unimodal cells in the PER can further dissociate different modality conditions

If unimodal neurons are invariably activated by a specific sensory input, their activity levels should remain constant between multimodal and their preferred unimodal conditions, reflecting the fact that both conditions contain the same image or sound of an object. However, it is also possible that unimodal cells are further modulated by different modality conditions while maintaining their preferred visual or auditory information. To examine the modulation of firing rates across modality conditions, we defined a rate modulation index (RMI) based on Cohen’s *d*, where larger *d* values indicate a greater difference between groups (see Methods). RMIs, calculated as the difference in mean firing rates between modality conditions, were determined for multimodal and visual conditions (VA – V) and multimodal and auditory conditions (VA – A).

Cells #10 and #11, examples of visual cells, are shown in **Figure 6A** with their RMIs. The subtracted value between multimodal and unimodal conditions (VA – V) was large and negative in both cells, indicating higher activities during the visual condition compared with the multimodal condition. Notably, visual cells exhibited “multisensory suppression”, such that firing rates were lower in the multimodal condition even though that condition contained the same visual information as the visual condition. However, VA – A values in both cells were small (near zero), indicating that their firing rates for multimodal conditions were not significantly different from those for auditory conditions. To visualize these patterns, we created scatter plots and histograms of RMI values for visual cells (**Fig. 6B**). VA – V values for visual cells were significantly different from zero (t_(20)_ = 8.9, *p* < 0.0001; one-sample t-test), indicating that visual cells further dissociated visual and multimodal conditions. However, VA – A values for visual cells were not significantly different from zero (t_(20)_ = 1.78, *p* = 0.091; one-sample t-test), suggesting that visual cells are not a suitable neuronal substrate for dissociating multimodal and auditory conditions.

**Fig. 6.**
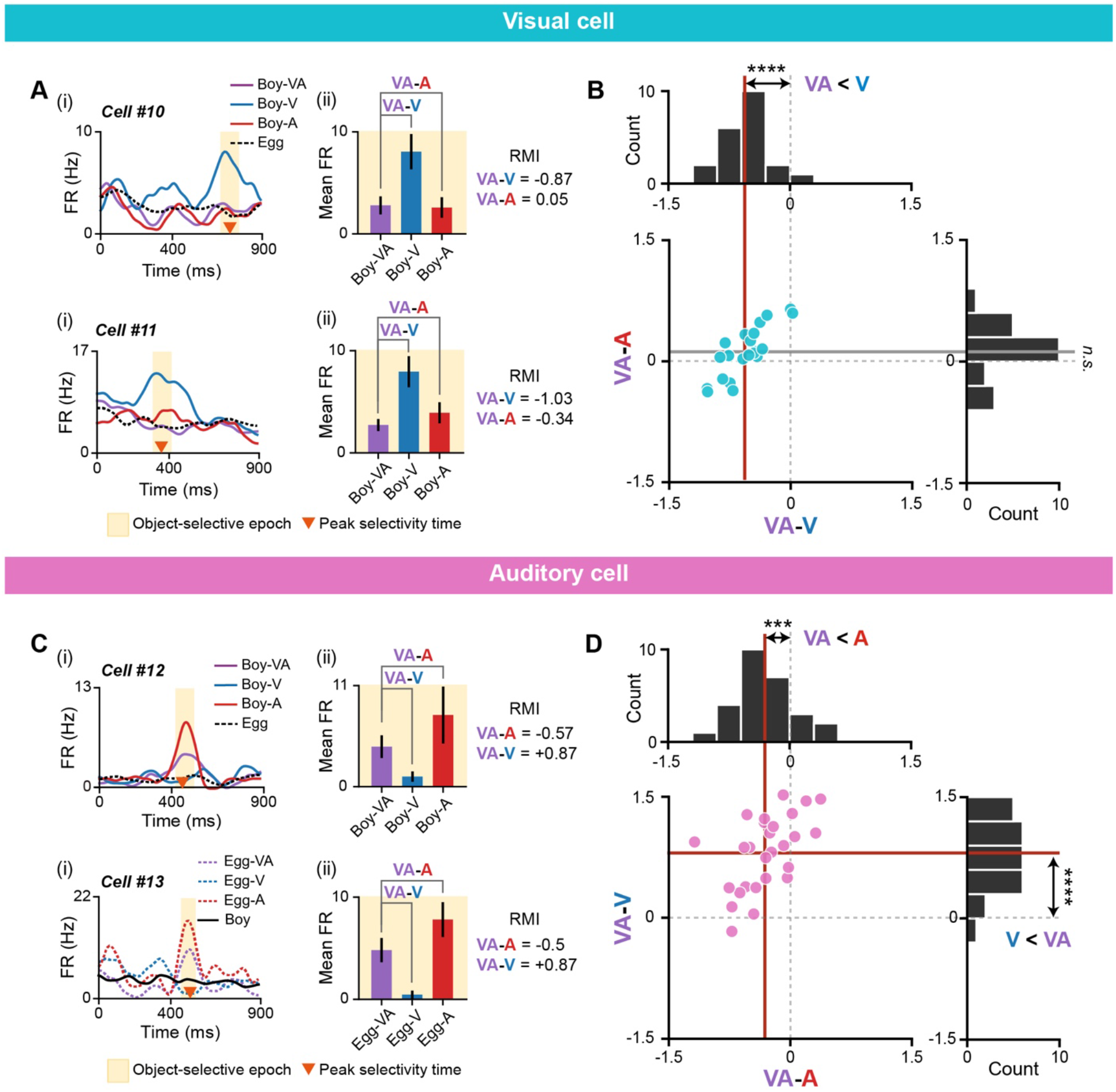
Further dissociation of modality conditions by visual and auditory cells. (**A**) Examples of visual cells (cells #10 and #11) demonstrating further dissociation of visual and multimodal conditions, but not multimodal and auditory conditions, as shown by their spike density functions (i) and mean firing rates within the selective epoch (ii). Differences in firing rate, quantified as RMI, showed that firing rates were different between visual and multimodal conditions (i.e., negative VA – V), but not between multimodal and auditory conditions (i.e., VA – A near zero). (**B**) Scatter plot and histograms of VA – V and VA – A in visual cells. For visual cells, the average VA – V (vertical red line) was significantly different from zero, whereas the average VA – A (horizontal gray line) was not. (**C**) Examples of auditory cells (cells #12 and #13) demonstrating further dissociation of all modality conditions, as shown by their spike density functions (i) and mean firing rates within the selective epoch (ii). RMIs showed that firing rates were different between auditory and multimodal conditions (i.e., negative VA – A), and also between multimodal and visual conditions (i.e., positive VA – V). (**D**) Scatter plot and histograms of VA – A and VA – V in auditory cells. The average VA – A (vertical red line) and average VA – V (horizontal red line) differed significantly from zero (\*\*\**p* < 0.001, \*\*\*\**p* < 0.0001; n.s., not significant).

Next, we examined RMI values in auditory cells (**Fig. 6C**). In cells #12 and #13, the mean firing rates for the auditory condition were higher than those in the multimodal condition (i.e., negative VA – A), although both conditions contained the same auditory information. That is, auditory cells, like visual cells, exhibited multisensory suppression. In addition, auditory cells further dissociated multimodal and visual conditions, showing relatively higher firing rates in the multimodal condition (i.e., positive VA – V). These patterns in auditory cells were visualized using scatter plots and histograms of RMI values (**Fig. 6D**). Further analyses showed that VA – A values for auditory cells were significantly different from zero (t_(26)_ = 4.48, *p* = 0.00013; one-sample t-test), indicating that these cells dissociated auditory and multimodal conditions. VA – V values for auditory cells were also significantly different from zero (t_(26)_ = 9.18, *p* < 0.0001; one-sample t-test).

Collectively, these findings demonstrate that visual, auditory, and multimodal conditions can be distinguished based on the firing rates of single auditory cells, which exhibited a rank order of firing rate of A > VA > V. Further analyses revealed that crossmodal cells exhibited heterogeneous patterns of neural modulation compared with unimodal cells (**Fig. S7**). The multisensory suppression displayed by both visual and auditory cells could not be explained by familiarity-coding for the multimodal condition (i.e., repetition suppression; **Fig. S8**). Taken together, these results suggest that unimodal cell types in the PER do not merely respond to the presence or absence of specific modality information. Instead, they are capable of differentially representing different modality conditions by modulating their firing rates according to the specific modality conditions.

### The PER neuronal population can decode object identities in both a modality-specific and modality-invariant manner

Having described different categories of object cells and their heterogeneous activity patterns in response to objects with different sensory modalities, we next sought to directly assess how PER neurons support multimodal object recognition. To this end, we conducted a population-decoding analysis using two different linear support vector machine (SVM) classifiers to evaluate distinct multimodal object-recognition processes. These two classifiers were designed to test whether the PER neurons as a population could decode object identities in a modality-specific manner (classifier 1; **Fig. 7A–C**) or a modality-invariant manner (classifier 2; **Fig. 7D– F**). For each classifier, we sought to determine if decoding performance was significant, and which cell categories contributed to the decoding.

**Fig. 7.**
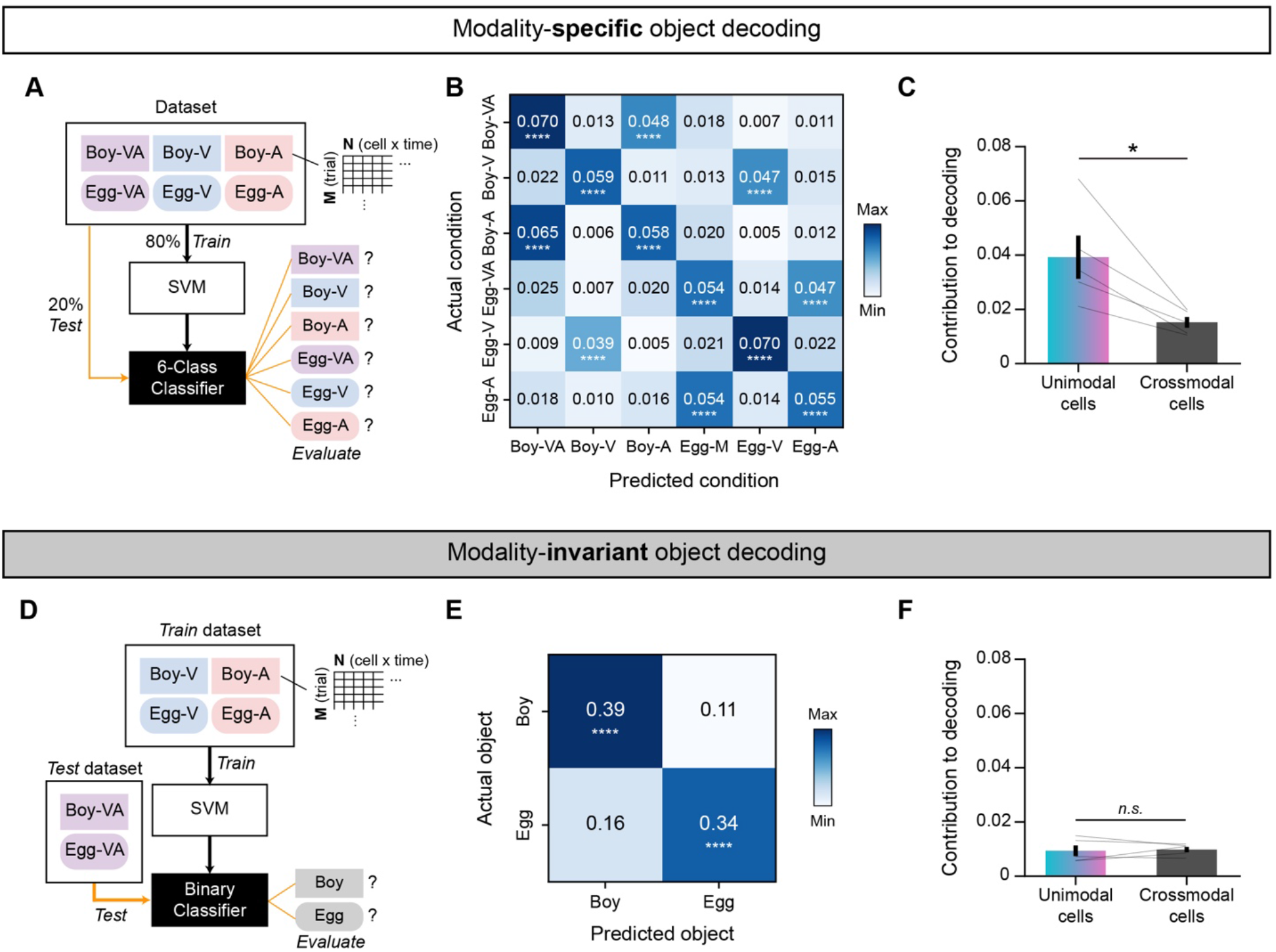
Modality-specific and modality-invariant decoding of object identities by the neuronal population in the PER. (**A**) Diagram summarizing modality-specific object decoding using a linear support vector machine (SVM). (**B**) Confusion matrix showing the average decoding accuracy of the classifier depicted in **A** (n = 5). (**C**) Comparison of the contribution of a single neuron to the decoding accuracy between unimodal and crossmodal cells, showing a significantly higher contribution of unimodal neurons to this type of decoding (n = 5). (**D**) Diagram summarizing the decoding of multimodal objects based on unimodal information (i.e., modality-invariant object decoding) with a linear SVM. Note that the classifier was trained with visual and auditory trials only, and tested on multimodal trials only. (**E**) Confusion matrix showing the average decoding accuracy of the classifier depicted in **E** (n = 5). (**F**) The contribution of a single neuron to modality-invariant object decoding was similar between unimodal and crossmodal cells. Data are presented as means ± SEM (\**p* < 0.05, \*\*\*\**p* < 0.0001; n.s., not significant).

For the first classifier, six object conditions – two objects, each with three modality conditions – were decoded using a 6-class SVM classifier (**Fig. 7A**). To create a dataset, we generated pseudo-populations of object cells and their firing rates during the task epoch for each rat (n = 5) by subsampling 5 trials from each condition (see Methods for details). We then employed a 5-fold cross-validation approach to train and test the dataset, repeating the procedures (subsampling, training, and testing) 100 times. A confusion matrix was created by averaging the proportions of actual and predicted conditions across rats (**Fig. 7B**). In the confusion matrix, the proportion in the diagonal line (i.e., decoding accuracy) was significantly higher compared with that in the shuffled distribution (*p* < 0.0001), indicating the successful decoding of both object identities and modality conditions (permutation test).

Next, we analyzed how unimodal and crossmodal cells, defined in the previous analysis (**Fig. 5E**), contributed to the decoding performance. We speculated that unimodal cells would make a greater contribution to the dissociation of modality conditions owing to their ability to dissociate not only visual and auditory inputs (**Fig. 5C**) but also unimodal and multimodal conditions (**Fig. 6B** and **6D**). For this analysis, we tested the respective contributions to decoding by quantifying the extent to which decoding accuracy decreased after shuffling data from a given cell category (see Methods for details). For example, to calculate the contribution of crossmodal cells to decoding, we shuffled trial labels (rows) only in features (columns) that were derived from crossmodal cells. We then assessed decoding accuracy before and after implementing this permutation, comparing the contribution of a single neuron in unimodal and crossmodal cell categories to decoding accuracy (**Fig. 7C**). Single unimodal cells exhibited significantly higher contributions to decoding accuracy compared with individual crossmodal neurons (t_(4)_ = 3.7, *p* = 0.021; paired t-test), indicating that the PER can decode modality-specific object information based on the activities of a limited number of unimodal cells.

Next, we investigated whether the neuronal population in the PER could achieve modality-invariant decoding of object identities. Specifically, we sought to determine if multimodal objects could be decoded solely from unimodal trials, by analogy to the ability of rats to retrieve multimodal objects when only unimodal cues are available (**Fig. 2B, 2C**, and **3D**). For this analysis, we trained the SVM to classify *Boy* and *Egg* objects using only unimodal trials (i.e., V and A). After training, we tested the classifier with multimodal trials to determine if the object identity could be successfully decoded (**Fig. 7D**). The creation of pseudo-populations followed a process similar to that described in the previous section. In the confusion matrix, the proportion along the diagonal, indicating the accuracy of invariant object decoding, was significantly higher than that in the shuffled distribution (*p* < 0.0001; permutation test) (**Fig. 7E**). Thus, successful modality-invariant decoding did not simply result from multimodal and unimodal conditions sharing the same choice response (**Fig. S9**).

Finally, we examined how different cell categories contributed to invariant object decoding (**Fig. 7F**). To measure the contribution to decoding, we quantified the degree of decrease in decoding accuracy after shuffling data from a given cell category (i.e., unimodal or crossmodal), as in **Figure 7C**. In contrast to the differentiation of modality information, the contribution of a single neuron to decoding performance was minimal for invariant objects. In addition, both crossmodal and unimodal cells contributed similarly to decoding (t_(4)_ = 0.29, *p* = 0.78; paired t-test) (**Fig. 7F**). These results suggest that the PER can also accomplish modality-invariant recognition of objects and further that this process is supported by population activity patterns of multiple neurons, rather than by a limited subset of single neurons.

## Discussion

In the current study, we investigated how the PER contributes to multimodal object recognition using a behavioral paradigm in which rats retrieved multimodal objects based on the objects’ multimodal or unimodal attributes. Rats identified multimodal objects correctly even when provided only unimodal cues, and the PER was required for normal performance. Single-unit recordings revealed that PER neurons exhibited transient object-selective signals that fired sequentially throughout the entire task epoch. Certain object-selective neurons responded primarily to visual or auditory attributes of an object (unimodal cells), whereas others exhibited equivalent selectivity across different object modalities (crossmodal cells). Unimodal cells further dissociated different modality conditions through modulation of their firing rates. Lastly, using a population-decoding analysis, we found that the PER was capable of accomplishing both modality-specific and modality-invariant object recognition. Specifically, modality-specific decoding was enabled by a small number of unimodal cells, whereas modality-invariant decoding was achieved through collective activity patterns of a relatively large number of neurons, regardless of their cell types. Overall, our findings suggest that the PER supports multimodal object recognition by engaging in both invariant recognition of a multimodal object and separation of object experiences based on modality information.

As previously reported, PER inactivation in our study resulted in performance deficits in the multimodal object-recognition task ^5,7^. Based on behavioral results, however, it remains uncertain whether the PER is important solely in “multimodal” situations. Specifically, because performance deficits were observed in both multimodal and unimodal conditions, the possibility remains that the role of the PER is limited to the separate processing of visual and auditory information ^6^. Indeed, it has been reported that the rodent PER is engaged in various tasks that employ visual-or auditory-only cues ^33,34^. A similar issue is applicable to previous behavioral experiments that reported performance deficits in tests of spontaneous object recognition in both crossmodal and unimodal conditions ^5–7^. Therefore, understanding the function of the PER in multisensory processing requires a detailed investigation of neural activity patterns under different modality conditions.

### Possible advantages of transient and sequential object selectivity within the PER

Since we controlled the sampling and response times of rats precisely by compelling nose-poke behaviors, we were able to describe the detailed temporal dynamics of neuronal activity during multimodal object recognition. We discovered that object-selective signals were elicited in PER neurons for a short period of time. However, this result is inconsistent with previous reports of persistent PER activity in both *in vitro* ^35,36^ and *in vivo* ^37^ settings. There are several possible explanations for why we did not observe persistent object selectivity. One possibility is that PER neurons in our study actually did maintain persistent firing, but object selectivity emerged transiently during the persistent firing. Most neurons analyzed in the current study were physiologically categorized as regular-spiking neurons, so their activities were rather persistent throughout the task epoch. In addition, it is important to note that the persistent selectivity of the PER reported in previous studies may be more closely related to neural correlates of a behavioral response than the stimulus itself. In our task, we were able to dissociate object- and response-selective signals by introducing a control condition. Notably, response signals displayed longer durations of selectivity compared with object selectivity (**Fig. S4**). We postulate that this long-lasting selectivity for the choice response might overlap with the previously reported persistent selectivity.

We also observed that object selectivity in the PER exhibited sequential characteristics. Although this sequential nature has rarely been observed in the PER, it is commonly reported in other brain regions, such as the prefrontal cortex ^38^, posterior parietal cortex ^39^, and hippocampus ^40^. This sequential pattern may have arisen because a specific behavioral sequence – maintaining nose-poke and then choosing left or right – was always evoked in our task. However, it should be noted that sequential coding has been reported to be beneficial for various aspects of memory processing. That is, a sequential activity pattern is a way to achieve high-dimensional information processing, which can enhance memory capacity and mitigate memory loss ^41^. It has also been suggested that sequential firing patterns within the medial temporal lobe represent temporal information of events, as exemplified by time cells in the hippocampus ^42^. The lateral entorhinal cortex, which receives extensive monosynaptic inputs from the PER, has also been reported to represent task-related time information ^43^. Thus, the PER may also contribute to the time component of episodic memory by representing both time and object information in an associative manner through sequential activity patterns.

### Operation of both integrated and segregated encoding of multimodal object information in the PER

Previous studies have described the PER as an associative area in terms of both its physiological characteristics ^29^ and task-related firing patterns ^26^. For example, neurons in the PER were found to be responsive to two paired visual stimuli that were associated with a reward outcome ^26^. The PER was also theorized to primarily function in the “unitization” process ^44^. That is, it was suggested that the PER plays a role in situations where complex features of a single entity must be integrated, such as when experiencing a complex object with multisensory information rather than sampling a simple cue. Based on these hypotheses, the PER is expected to encode multimodal objects in an integrated fashion instead of representing information of a single object separately based on its modality. Consistent with these expectations, we discovered that most object cells in the PER exhibit constant selectivity patterns, irrespective of the modality condition (i.e., crossmodal cells). We believe that our task requirements were suitable for facilitating the unitization process, as the multisensory cues were spatially and temporally congruent, and each audiovisual combination required the same behavioral response. Thus, our results provide experimental support for the idea that single neurons in the PER can encode multimodal objects in a unitized representation.

However, it should also be noted that a significant proportion of unimodal cells in the PER primarily responded to a specific sensory modality when processing object information, an outcome that is not expected based on previous literature reports ^31,44^. These neurons not only preferred a particular sensory modality, they also further dissociated unimodal and multimodal conditions through modulation of their firing rates. These unimodal activities could be interpreted as purely perceptual signals that reflect the physical attributes of visual and auditory cues. The perceptual-mnemonic hypothesis, which posits that the PER is involved in both perception and memory, may further support the interpretation that unimodal cells indeed represent perceptual information ^45–49^. However, it is unlikely that unimodal neurons simply mirrored low-level perceptual features of the stimuli. If unimodal cells represented perceptual signals originating from the visual (or auditory) cortex, it is likely that the posterior (or anterior) PER would have more visual (or auditory) cells since visual (or auditory) input is more dominant in the corresponding area. Instead, we observed that each cell category appeared to be equally distributed along the anteroposterior axis of the PER. Moreover, unimodal cells showed modulation by their non-preferred sensory modality, indicating that they were not simply responding to the presence of a specific modality cue. Thus, unimodal cell activity in this area could have been driven by intrinsic connections within the PER ^29^ or by inputs from other higher-order associative areas, such as the prefrontal cortex and hippocampus ^50–52^. Given that the PER is part of the medial temporal lobe memory system, it can be argued that unimodal representations exist for memory encoding and retrieval rather than for simple sensory processing.

### Dual functions of the PER in multimodal object recognition: invariant recognition and episodic memory

From a computational standpoint, an object-recognition system should be able to recognize an object through an invariant representation, even if the object’s physical attributes are modified ^12^. In multimodal object recognition, it is also important that objects be identified invariantly to modality information. This modality invariance can be attained by individual neurons, as exemplified by “concept cells” that fire invariantly to both the image and voice of a person ^53–55^. Crossmodal cells in our study shared some commonalities with concept cells from the human hippocampus as they showed some degree of invariance to modality information when coding object identities. However, we discovered that individual crossmodal cells within the PER do not contribute significantly to modality-invariant object recognition, making contributions to decoding accuracy similar to those of the unimodal cell type. This may be because crossmodal cells were not fully invariant to modality conditions, but instead showed slight modulations in response to different modality conditions of objects (**Fig. S7**). More detailed investigations of concept-like representations also suggest that firing patterns of individual neurons can be heterogeneous, and that population-level activities are better suited to achieve invariance ^56,57^.

In addition to the invariant recognition process, we discovered that populations of PER neurons can perform modality-specific object decoding, a process that seems to be counterproductive for the invariant identification of objects. However, in terms of episodic memory, segregation of similar events (i.e., pattern separation) is a crucial computational step for encoding and retrieving correct memory ^58,59^. In cases where a single object is experienced by multiple senses, each experience should be separated into different episodes, even though they involve the same object. Pattern separation for episodic memory is thought to be primarily implemented in the dentate gyrus ^60,61^. However, since a significant portion of information received by the dentate gyrus relies on connections between the PER and entorhinal cortex, modality-specific information in the PER could be an essential source for pattern separation within the dentate gyrus. In addition, it has been suggested that the PER itself can support pattern separation when two visual stimuli are highly overlapped as they morph into each other ^49^. Validating the relationship between modality-specific representations and pattern separation will require future studies that systematically manipulate the amount of information from each modality.

## Methods

### Subjects

Male Long-Evans rats (10 wk old; n = 14) were obtained and individually housed in a temperature- and humidity-controlled animal colony. Rats were allowed free access to food and water for 1 wk before food restriction, during which they were allowed only 2 to 3 pellets (6–10 g) per day to maintain them at ∼80% of their free-feeding body weight (∼400–420 g). Rats were housed on a 12-h light/dark cycle (lights on at 8 AM), and all experiments were performed in the light phase. All animal procedures were performed in accordance with the regulations of the International Animal Care and Use Committee of Seoul National University.

### Behavioral apparatus

The apparatus consisted of an elevated chamber (22 × 35 × 40 cm; 94 cm above the floor) with a custom-built device (22 × 18 cm) at the front of the chamber that was used for manipulating cues and measuring animal behaviors with Arduino MEGA (Arduino) and MATLAB (MathWorks). The frame of the device was printed with a 3D printer (Mojo; Stratasys), and the center of the device contained a transparent acrylic window (8 × 10 cm) with a nose-poke hole (diameter, 2.4 cm; depth, 1.5 cm). The hole was equipped with an infrared sensor for measuring the onset and maintenance of nose-poking behaviors during cue sampling. An LCD panel (3.5 inch; Nextion) for presenting a visual cue, operated by Arduino, was positioned behind the acrylic window. Directly behind the LCD panel was a 3W speaker, operated through an Arduino music player module (DFPlayer Mini Mp3 Player; DFRobot), for presenting an auditory cue. The device contained two identical ports located on the left and right side. Each port was equipped with a servo-motorized door for controlling access and infrared sensors for detecting choice responses. Another servo-motorized door located on the top of the port controlled the gravity-fed delivery of a pre-loaded food reward to the choice port. A small buzzer was placed on the back of the chamber to provide auditory feedback about the correctness of the rat’s choice. The experimental room was dimly lit with a circular array of LEDs (0.8 lux), and white noise (68 dB) was played through loudspeakers to block out uncontrolled noise.

### Behavioral paradigm

#### Shaping

After 6 d of handling, a shaping stage was included during which rats learned how to maintain nose-poking of the center hole. The required duration for nose-poke was 10 ms beginning in the first shaping trial, and then was increased by 10 ms for each successful poke to a maximum of 400 ms. When rats failed to maintain their nose-poke for the required duration, the trial was stopped and a 4-s interval was given together with auditory feedback (buzzer, 230 Hz, 76 dB). Once rats successfully completed 100 trials of 400-ms nose-pokes within a 30-min session, they advanced to the multimodal object-recognition task.

#### Multimodal object recognition – training

Rats learned to make an associated choice response based on a presented cue. Initially, the rats were trained under multimodal object conditions (designated VA), in which a combination of visual and auditory cues was presented simultaneously. The visual cues used were 2D photographic images of two junk objects – a boy and an egg – presented via an LCD panel (1.6 lux). The two object images were adjusted to equal luminance by matching their average gray values in Photoshop (Adobe). Auditory cues were 5 kHz and 10 kHz sine-wave tones (81 dB) that briefly repeated twice. Each object was associated with either a left or right choice response. The combination of audiovisual cue and stimulus-response contingency was counterbalanced across rats. An object containing a boy (or egg) image was called a *Boy* (or *Egg*) object, regardless of the auditory cue associated with it. Nose-poking to the center hole simultaneously triggered the onset of visual and auditory cues, which remained presented for up to 400 ms while the rat maintained the nose-poke. If rats failed to maintain the nose-poke for at least 400 ms (i.e., prematurely withdrawn nose-poke), cues disappeared and the auditory feedback was given together with a 4-s interval. On the next nose-poking, a pseudo-random stimulus was presented regardless of the previously experienced stimulus. Prematurely withdrawn nose-pokes did not increase trial numbers. In successful nose-pokes (>400 ms), the doors covering the left and right choice ports were opened, allowing the rat to access one of the choice ports. A correct choice response resulted in delivery of a food reward, whereas incorrect responses resulted in auditory feedback without a food reward together with an 8-s inter-trial interval. Rats performed 100 to 120 trials in total within a session. After rats exceeded the learning criterion (>75% correct in all conditions for two consecutive days), they learned the same task but using two simple visual cues as a control (C) condition. Rats that exceeded the learning criterion in the control condition were then trained with both multimodal objects and control stimuli within a session until they reached the criterion. After completing all training procedures, rats underwent either cannula or hyperdrive implantation surgery (see below for details). After surgery, they were again trained simultaneously on multimodal and control conditions and then proceeded to the test phase.

#### Multimodal object recognition – testing

Unimodal conditions (visual or auditory) were introduced for the first time in the test phase of multimodal object recognition. In the visual (V) condition, only the boy or egg image was presented without an auditory cue. In the auditory (A) condition, only a 5 or 10 kHz sound was presented without an image. Rats were required to make the same choice response associated with the multimodal object based on the unimodal stimulus. In the drug-infusion study, rats were serially tested under multimodal, visual, auditory, and control conditions in separate sessions and performed 120 trials per session. In the electrophysiological study, all eight conditions (two objects × three modality conditions plus two control stimuli) were pseudo-randomly presented within a session, and rats performed 180 to 240 trials per session (see below for details).

### Drug infusion

The guide cannula (24 gauge, 18 mm long), internal cannula (30 gauge, 19 mm long), and dummy cannula (30 gauge, 19 mm long) were built in-house. A surgery targeting the bilateral PER was performed by first carefully retracting the left and right temporalis muscle, after which two holes were drilled bilaterally on the skull surface (4.8 mm posterior to bregma, 5.2 mm lateral to the midline). Guide cannulas were angled 15 degrees outward, lowered to 7 mm below the cortical surface, and chronically fixed with four anchoring screws and dental cement. The procedure was completed by placing dummy cannulas inside the guide cannulas. During insertion, the tips of internal and dummy cannulas were protruded 1 mm from the tip of guide cannulas. Cannulas were cleaned at least once every 2 d. The drug infusion schedule was started after all rats had been retrained to multimodal and control conditions. PBS (0.5 μl per site) and the GABA-A receptor antagonist, muscimol (MUS; 0.5 μl per site), were bilaterally injected into the PER on alternate days using a Hamilton syringe (10 μl). After one rat (rat #5) showed immobilization side effects following muscimol injection, the injection amount was reduced to 0.3 μl. Drug infusions were made 20 min before the start of the behavioral experiment. Rats were tested in each condition on a different day in the following order: multimodal, unimodal (visual and auditory), and control. The order of visual and auditory sessions was pseudo-randomized for each rat. At the end of the experiment (20 min before sacrifice), the diffusion range of MUS was estimated by injecting rats with fluorescent BODIPY TMR-X–labeled MUS (fMUS) and monitoring fMUS by fluorescence microscopy.

### Hyperdrive implantation

The hyperdrive containing 27 tetrodes was built in-house. Tetrodes were prepared by winding together four formvar-insulated nichrome wires (diameter, 17.8 µm) and bonding them with heat. Impedance was reduced to ∼200 kΩ at 1 kHz by gold-plating wires using a Nano-Z plating system (Neuralynx). For targeting the PER along the anteroposterior axis, a 12G stainless-steel cannula bundle housing 27 tetrodes was formed into an elliptical shape (major axis, 3.4–3.8 mm; minor axis, 2–2.4 mm). After performing surgery to target the right hemisphere of the PER, as described above, a hole sized to fit the tetrode bundle was drilled on the skull surface. The bundle tip was angled 12 degrees outward and lowered until it touched the cortical surface, after which the hyperdrive was chronically fixed with 11 anchoring screws and bone cement.

### Electrophysiological recording

After allowing 3 d to recover from surgery, rats were reacclimated to experimentation by handling for 4 d and then retrained to perform the multimodal object recognition task under multimodal and control conditions. Individual tetrodes were lowered daily. After most of the tetrodes had reached the PER and rats showed greater than 75% correct responses in both multimodal and control conditions for two consecutive days, recording sessions were begun. In the recording sessions, the unimodal condition was introduced for the first time, such that multimodal, visual, auditory, and control conditions were all presented pseudo-randomly during a session. Recordings were conducted in each rat for 5 to 6 d, and no attempt was made to record the same neuron across days. Neural signals were amplified 1000–10,000-fold and bandpass filtered (300–6000 Hz) using a Digital Lynx data-acquisition system (Neuralynx). Spike waveforms exceeding a preset threshold (adjusted within the range of 40–150 µV) were digitized at 32 kHz and timestamped.

### Histology

Rats were sacrificed with an overdose of CO_2_ and transcardially perfused first with PBS and then with a 4% (v/v) formaldehyde solution. The brain was extracted and maintained in a 4% (v/v) formaldehyde-30% sucrose solution at 4°C until it sank to the bottom of the container. The brain was subsequently coated with gelatin, soaked again in 4% (v/v) formaldehyde-30% sucrose solution, and then sectioned at a thickness of 40 μm using a freezing microtome (HM 430; ThermoFisher Scientific). For every three consecutive sections, the second and third sections were mounted for staining. For the drug infusion study (n = 6), every second section was Nissl-stained with thionin solution, and every third section was stained with DAPI solution (Vectashield) for fluorescence microscopy. For the electrophysiological study (n = 8), every second section was stained with thionin solution, and every third section was stained with gold solution for myelin staining. Photomicrographs of each brain section were obtained using a microscope mounted with a digital camera (Eclipse 80i; Nikon). To accurately estimate the position of tetrodes, we reconstructed the configuration of tetrodes based on histology results, and then compared it with the actual configuration to match the numbering of the tetrodes (Voxwin, UK).

### Unit isolation

All single units were manually isolated using a custom program (WinClust), as previously described ^32,62^. Various waveform parameters (i.e., peak amplitude, energy, and peak-to-trough latencies) were used for isolating single units, but peak amplitude was the primary criterion. Units were excluded if more than 1% of spikes occurred within the refractory period (1 ms) and mean firing rates during the task epoch (from cue onset to response) were lower than 0.5 Hz.

### Single-unit analysis

#### Basic firing properties

Single units were grouped into bursting, regular-spiking, and unclassified neurons based on their autocorrelograms and interspike-interval histograms (Bartho et al., 2004). Specifically, cells were classified as bursting neurons if they met the following criterion:

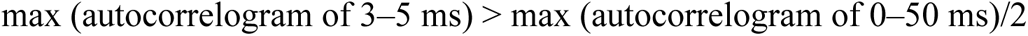

Among the remaining neurons, those in which the mode of the interspike-interval histogram was less than 35 ms were classified as regular-spiking neurons. Neurons that did not belong to either group were categorized as unclassified neurons. Spike width was measured as the distance from peak to trough.

#### Trial filtering

All subsequent analyses described below were performed using correct trials only. An overview of the subsequent single-unit analysis process is presented in **Figure S3**. To control for variability in response latency (i.e., from cue offset to the end of choice response), we excluded trials where the latency exceeded 3 absolute median deviations of all correct trials. If a recording session had less than five correct trials in any of the eight stimulus conditions, all units recorded in that session were excluded from further analysis.

#### Defining selective epoch

Firing rates were calculated within 50-ms time bins with increments of 10 ms. All subsequent analyses described below were performed on firing rates within the task epoch, defined as the 900-ms interval from the start of the sample phase to immediately preceding the end of the response phase. To identify a selective epoch in which firing rates were significantly different between *Boy* and *Egg* objects, we performed two-way repeated measures ANOVA (object identity and modality condition as two factors) in each time bin using trials from object conditions (two objects with three modality conditions). The time bin with the largest effect size (η^2^) for the object identity factor was designated “peak selectivity time”, representing the moment when the firing rate difference between the two objects was maximal. The selective epoch was defined as having more than five consecutive time bins around the peak selectivity time, each with a p-value < 0.05 for the object identity factor.

#### Multiple linear regression

The following multiple linear regression models were used to describe firing patterns in relation to task-related conditions:

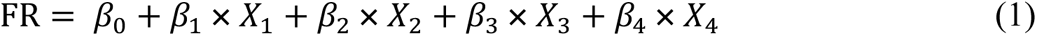

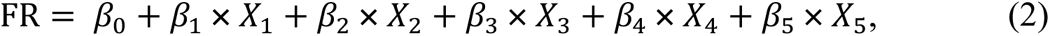

where the dependent variable FR, is the firing rate within the selective epoch, described above. In the standard model (1), β_1_ is the constant term, β_1_ × *X*_1_ is the term for visual information of the preferred object, β_2_ × *X*_2_ is the term for auditory information of the preferred object, β_3_ × *X*_3_ is the term for visual information of the non-preferred object, and β_4_ × *X*_4_ is the term for auditory information of the non-preferred object. The independent variables (*X*) were binary coded to reflect the existence of an image or sound for an object. For example, if a neuron was classified as a *Boy*-preferring object cell, *X*_1_ had a value of one in Boy-VA and Boy-V trials, and zero in all other conditions. In the extended model (2), the term β_5_ × *X*_5_ was added to further examine the influence of the response factor. X_5_ had a value of one if a trial required a left choice response, and zero if it required a right choice response. All trial conditions (VA, V, A, C) were used to estimate the regression model. β coefficients were standardized by z-scoring both dependent and independent variables prior to regression fitting. To dissociate neurons that were mainly modulated by choice responses (i.e., response cell) rather than object information, we quantified how much the model was improved by adding the response factor. Specifically, we subtracted the AIC (Akaike Information Criterion) for the extended model (2) from that for the standard model (1). If a neuron exhibited a significantly higher AIC difference, we concluded that most of its activity patterns were explained by the response factor, and thus classified it as a response cell. The significance of the AIC difference was determined by comparison with the null distribution, obtained by shuffling trial conditions (shuffled 1000 times; α = 0.01). Neurons with a selective epoch but not classified as response cells were categorized as object cells. To describe how object cells responded to different modality information, we examined regression coefficients in the standard model (1) using β_1_ and β_2_ to quantify how strongly an object cell responded to visual and auditory information, respectively, of a preferred object. We did not further examine regression coefficients for a non-preferred object (i.e., β_3_ and β_4_) (see **Fig. S6**). Neurons for which the difference between β_1_ and β_2_ was significantly higher or lower than the difference obtained after shuffling trial conditions were classified as visual or auditory cells, respectively (shuffled 1000 times; α = 0.05, two-sided permutation test).

#### Rate modulation index

We calculated a “rate modulation index” (RMI) to quantify increases or decreases in a neuron’s firing rates in the multimodal condition relative to the unimodal condition. Firing rate differences between the multimodal and unimodal condition were quantified using Cohen’s *d* as follows:

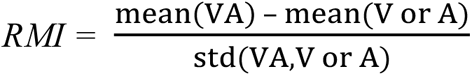

The index was calculated only in the modality conditions of the preferred object, and was referred to as “VA – V” when the index was calculated between multimodal and visual conditions, and "VA – A" when it was calculated between multimodal and auditory conditions.

### Population decoding

A linear support vector machine (*sklearn.svm.SVC,* Python function), with cost parameter set to 0.01, was used for population decoding. Population decoding was performed on rats in which at least 20 object cells were recorded across sessions (5 of 8 rats). Spikes were binned into 100-ms time bins within the task epoch (900-ms duration) and z-scored. Pseudo-populations of neurons were constructed in each rat as follows: For each object cell, five trials for each of the six object conditions (two objects × three modalities) were subsampled. Firing rates in the subsampled trials were horizontally concatenated to the pseudo-population. Thus, each pseudo-population had 30 rows (5 trials × 6 conditions) and N columns (or features), where N was the number of time bins (9) multiplied by the number of object cells. For modality-specific object decoding (**Fig. 7A**), the entire subsampled dataset (30 samples) was used for both training and testing.

One-vs.-one classification was performed using stratified 5-fold cross-validation. For modality-invariant object decoding (**Fig. 7D**), a binary classifier was trained using only unimodal trials, and then tested with multimodal trials ^63^. We did not perform cross-validation here since the training and test sets were completely separate. Subsampling, training, and testing were repeated 100 times in both decoding procedures, and the average of these repeated results was used as the representative value for each rat. A permutation test, performed by shuffling trial conditions, was used for significance testing (shuffled 1000 times; α = 0.05). Confusion matrices (**Fig. 7B** and **7E**) were constructed by averaging the results from all rats. Contributions to decoding performance (**Fig. 7C** and **7F**) were measured using the permutation feature importance method. Specifically, after training the classifier, we selected all features from a given cell category (unimodal or crossmodal) and shuffled their rows (or trial labels) to break the relationship between the true label and selected features. The decrease in decoding accuracy after permutation was used as an indicator of how much the selected features contributed to decoding performance. Contribution to decoding was calculated as follows:

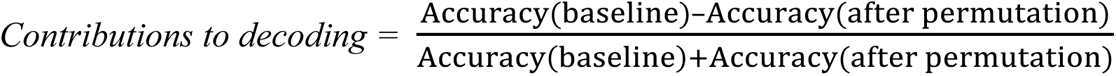

To measure the contribution of a single cell to decoding performance in a given category, we divided the value by the number of cells in that category within each rat.

### Quantification and statistical analysis

Data were statistically tested using custom-made codes written in MATLAB and Python. Student’s t-test, analysis of variance (ANOVA), Wilcoxon sign-rank test, Chi-square test, and permutation test were used for statistical comparisons. A one-sample t-test was used to verify that the behavioral performance was above the level of chance and RMI values were significantly different from zero. One-way repeated measures ANOVA was implemented for comparing behavioral results across modality conditions. Two-way repeated measures ANOVA was used to compare behavioral results (drug and modality condition as two factors), as well as to identify object-selective epoch (object and modality condition as two factors). Post hoc analyses were carried out using t-test with p-values corrected using the Holm-Bonferroni method. Wilcoxon signed-rank test was used to compare the regression coefficients, β_1_ and β_2_. An ordinary least squares method was used for both multiple and simple linear regression. Chi-square test was used for comparisons of proportions. A permutation test was used for categorizing response-selective neurons and defining significance levels for population decoding accuracy. Unless otherwise indicated, the significance level was set at α = 0.05. Error bars indicate standard error of the mean (SEM) unless stated otherwise.

## Author contributions

Conceptualization, H.-Y.L. and I.L.; Methodology, H.-Y.L. and I.L.; Software, H.-Y.L. and I.L.; Validation, H.-Y.L. and I.L.; Formal analysis, H.-Y.L.; Investigation, H.-Y.L.; Resources, I.L.; Data curation, H.-Y.L. and I.L.; Wiring – Original Draft, H.-Y.L.; Wiring – Review & Editing, H.-Y.L. and I.L.; Visualization, H.-Y.L. and I.L.; Supervision, I.L.; Project Administration, I.L.; Funding Acquisition, I.L.

## Data Availability

The datasets generated and/or analyzed during the current study are available from the corresponding author upon reasonable request.

## Acknowledgments

This study was supported by basic research grants (NRF - 2019R1A2C2088799, 2021R1A4A2001803, 2022M3E5E8017723) from the National Research Foundation of Korea and the BK21 program.

## References

1. Jones, E.G., and Powell, T.P.S. (1970). An Anatomical Study of Converging Sensory Pathways. Brain 93, 793–820. 10.1037/a0015325.

2. Ghazanfar, A.A., and Schroeder, C.E. (2006). Is neocortex essentially multisensory? Trends in Cognitive Sciences 10, 278–285. 10.1016/j.tics.2006.04.008.

3. Davenport, R.K., and Rogers, C.M. (1970). Intermodal Equivalence of Stimuli in Apes. Science 168, 279–280. 10.1126/science.168.3928.279.

4. Murray, E.A., and Mishkin, M. (1985). Amygdalectomy Impairs Crossmodal Association in Monkeys. Science 228, 604–606. 10.1126/science.3983648.

5. Winters, B.D., and Reid, J.M. (2010). A Distributed Cortical Representation Underlies Crossmodal Object Recognition in Rats. Journal of Neuroscience 30, 6253–6261. 10.1523/JNEUROSCI.6073-09.2010.

6. Albasser, M.M., Amin, E., Iordanova, M.D., Brown, M.W., Pearce, J.M., and Aggleton, J.P. (2011). Separate but interacting recognition memory systems for different senses: The role of the rat perirhinal cortex. Learning & Memory 18, 435–443. 10.1101/lm.2132911.

7. Jacklin, D.L., Cloke, J.M., Potvin, A., Garrett, I., and Winters, B.D. (2016). The Dynamic Multisensory Engram: Neural Circuitry Underlying Crossmodal Object Recognition in Rats Changes with the Nature of Object Experience. Journal of Neuroscience 36, 1273–1289. 10.1523/JNEUROSCI.3043-15.2016.

8. Bruck, J.N., Walmsley, S.F., and Janik, V.M. (2022). Cross-modal perception of identity by sound and taste in bottlenose dolphins. Science Advances 8, eabm7684. 10.1126/sciadv.abm7684.

9. Solvi, C., Al-Khudhairy, S.G., and Chittka, L. (2020). Bumble bees display cross-modal object recognition between visual and tactile senses. Science 367, 910–912. 10.1126/science.aay8064.

10. Ito, M., Tamura, H., Fujita, I., and Tanaka, K. (1995). Size and position invariance of neuronal responses in monkey inferotemporal cortex. Journal of Neurophysiology 73, 218–226. 10.1152/jn.1995.73.1.218.

11. Booth, M.C., and Rolls, E.T. (1998). View-invariant representations of familiar objects by neurons in the inferior temporal visual cortex. Cerebral Cortex 8, 510–523. 10.1093/cercor/8.6.510.

12. DiCarlo, J.J., Zoccolan, D., and Rust, N.C. (2012). How does the brain solve visual object recognition? Neuron 73, 415–434. 10.1016/j.neuron.2012.01.010.

13. Mishkin, M. (1978). Memory in monkeys severely impaired by combined but not by separate removal of amygdala and hippocampus. Nature 273, 297–298. 10.1038/273297a0.

14. Ennaceur, A. (2010). One-trial object recognition in rats and mice: Methodological and theoretical issues. Behavioural Brain Research 215, 244–254. 10.1016/j.bbr.2009.12.036.

15. Fahy, F.L., Riches, I.P., and Brown, M.W. (1993). Neuronal activity related to visual recognition memory: long-term memory and the encoding of recency and familiarity information in the primate anterior and medial inferior temporal and rhinal cortex. Exp Brain Res 96, 457–472. 10.1007/BF00234113.

16. Ahn, J.-R., and Lee, I. (2015). Neural Correlates of Object-Associated Choice Behavior in the Perirhinal Cortex of Rats. Journal of Neuroscience 35, 1692–1705. 10.1523/JNEUROSCI.3160-14.2015.

17. Zola-Morgan, S., Squire, L.R., Amaral, D.G., and Suzuki, W.A. (1989). Lesions of perirhinal and parahippocampal cortex that spare the amygdala and hippocampal formation produce severe memory impairment. The Journal of neuroscience : the official journal of the Society for Neuroscience 9, 4355–4370. 10.1523/JNEUROSCI.09-12-04355.1989.

18. Burke, S.N., Maurer, A.P., Hartzell, A.L., Nematollahi, S., Uprety, A., Wallace, J.L., and Barnes, C.A. (2012). Representation of three-dimensional objects by the rat perirhinal cortex. Hippocampus 22, 2032–2044. 10.1002/hipo.22060.

19. Deshmukh, S.S., Johnson, J.L., and Knierim, J.J. (2012). Perirhinal cortex represents nonspatial, but not spatial, information in rats foraging in the presence of objects: Comparison with lateral entorhinal cortex. Hippocampus 22, 2045–2058. 10.1002/hipo.22046.

20. Norman, G., and Eacott, M.J. (2005). Dissociable effects of lesions to the perirhinal cortex and the postrhinal cortex on memory for context and objects in rats. Behavioral Neuroscience 119, 557–566. 10.1037/0735-7044.119.2.557.

21. Suzuki, W.A., and Amaral, D.G. (1994). Perirhinal and parahippocampal cortices of the macaque monkey: Cortical afferents. Journal of Comparative Neurology 350, 497–533. 10.1002/cne.903500402.

22. Burwell, R.D., and Amaral, D.G. (1998). Cortical afferents of the perirhinal, postrhinal, and entorhinal cortices of the rat. Journal of Comparative Neurology 398, 179–205. 10.1002/(SICI)1096-9861(19980824)398:2<179::AID-CNE3>3.0.CO;2-Y.

23. Burwell, R.D. (2006). The Parahippocampal Region: Corticocortical Connectivity. Annals of the New York Academy of Sciences 911, 25–42. 10.1111/j.1749-6632.2000.tb06717.x.

24. Taylor, K.I., Moss, H.E., Stamatakis, E.A., and Tyler, L.K. (2006). Binding crossmodal object features in perirhinal cortex. Proceedings of the National Academy of Sciences 103, 8239–8244. 10.1073/pnas.0509704103.

25. Holdstock, J.S., Hocking, J., Notley, P., Devlin, J.T., and Price, C.J. (2009). Integrating visual and tactile information in the perirhinal cortex. Cerebral Cortex 19, 2993–3000. 10.1093/cercor/bhp073.

26. Sakai, K., and Miyashita, Y. (1991). Neural organization for the long-term memory of paired associates. Nature 354, 152–155. 10.1038/354152a0.

27. Naya, Y., Yoshida, M., and Miyashita, Y. (2003). Forward processing of long-term associative memory in monkey inferotemporal cortex. The Journal of neuroscience : the official journal of the Society for Neuroscience 23, 2861–2871. 23/7/2861 [pii].

28. Suzuki, W. a., and Naya, Y. (2014). The Perirhinal Cortex. Annual Review of Neuroscience 37, 39–53. 10.1146/annurev-neuro-071013-014207.

29. Unal, G., Apergis-Schoute, J., and Paré, D. (2012). Associative Properties of the Perirhinal Network. Cerebral Cortex 22, 1318–1332. 10.1093/cercor/bhr212.

30. Ohnuki, T., Osako, Y., Manabe, H., Sakurai, Y., and Hirokawa, J. (2020). Dynamic coordination of the perirhinal cortical neurons supports coherent representations between task epochs. Communications biology, 1-. 10.1038/s42003-020-01129-3.

31. Fiorilli, J., Marchesi, P., Ruikes, T., Veld, G.H. in ‘t, Buckton, R., Quintero, M.D., Reiten, I., Bjaalie, J., and Pennartz, C.M.A. (2023). Neural correlates of object identity and reward outcome in the corticohippocampal hierarchy: double dissociation between perirhinal and secondary visual cortex. Preprint at bioRxiv, 10.1101/2023.05.24.542117 10.1101/2023.05.24.542117.

32. Lim, H., Ahn, J., and Lee, I. (2022). The Interaction of Cue Type and Its Associated Behavioral Response Dissociates the Neural Activity between the Perirhinal and Postrhinal Cortices. eNeuro 9, 1–17.

33. Park, E.H., Ahn, J.-R., and Lee, I. (2017). Interactions between stimulus and response types are more strongly represented in the entorhinal cortex than in its upstream regions in rats. eLife 6, 1–11. 10.7554/eLife.33415.001.

34. Kholodar-Smith, D.B., Boguszewski, P., and Brown, T.H. (2008). Auditory trace fear conditioning requires perirhinal cortex. Neurobiology of Learning and Memory 90, 537–543. 10.1016/j.nlm.2008.06.006.

35. Beggs, J.M., Moyer, J.R., McGann, J.P., and Brown, T.H. (2000). Prolonged Synaptic Integration in Perirhinal Cortical Neurons. Journal of Neurophysiology 83, 3294–3298. 10.1152/jn.2000.83.6.3294.

36. Navaroli, V.L., Zhao, Y., Boguszewski, P., and Brown, T.H. (2012). Muscarinic receptor activation enables persistent firing in pyramidal neurons from superficial layers of dorsal perirhinal cortex. Hippocampus 22, 1392–1404. 10.1002/hipo.20975.

37. Bos, J.J., Vinck, M., van Mourik-Donga, L.A., Jackson, J.C., Witter, M.P., and Pennartz, C.M.A. (2017). Perirhinal firing patterns are sustained across large spatial segments of the task environment. Nature Communications 8, 15602. 10.1038/ncomms15602.

38. Bolkan, S.S., Stujenske, J.M., Parnaudeau, S., Spellman, T.J., Rauffenbart, C., Abbas, A.I., Harris, A.Z., Gordon, J.A., and Kellendonk, C. (2017). Thalamic projections sustain prefrontal activity during working memory maintenance. Nature Neuroscience 20, 987–996. 10.1038/nn.4568.

39. Harvey, C.D., Coen, P., and Tank, D.W. (2012). Choice-specific sequences in parietal cortex during a virtual-navigation decision task. Nature 484, 62–68. 10.1038/nature10918.

40. Pastalkova, E., Itskov, V., Amarasingham, A., and Buzsáki, G. (2008). Internally Generated Cell Assembly Sequences in the Rat Hippocampus. Science 321, 1322–1327. 10.1126/science.1159775.

41. Rajan, K., Harvey, C.D., and Tank, D.W. (2016). Recurrent Network Models of Sequence Generation and Memory. Neuron 90, 128–142. 10.1016/j.neuron.2016.02.009.

42. Kraus, B.J., Robinson, R.J., White, J.A., Eichenbaum, H., and Hasselmo, M.E. (2013). Hippocampal “Time Cells”: Time versus Path Integration. Neuron 78, 1090–1101. 10.1016/j.neuron.2013.04.015.

43. Tsao, A., Sugar, J., Lu, L., Wang, C., Knierim, J.J., Moser, M.-B., and Moser, E.I. (2018). Integrating time from experience in the lateral entorhinal cortex. Nature 561, 57–62. 10.1038/s41586-018-0459-6.

44. Fiorilli, J., Bos, J.J., Grande, X., Lim, J., Düzel, E., and Pennartz, C.M.A. (2021). Reconciling the object and spatial processing views of the perirhinal cortex through task-relevant unitization. Hippocampus, 1–19. 10.1002/hipo.23304.

45. Eacott, M.J., Gaffan, D., and Murray, E.A. (1994). Preserved Recognition Memory for Small Sets, and Impaired Stimulus Identification for Large Sets, Following Rhinal Cortex Ablations in Monkeys. European Journal of Neuroscience 6, 1466–1478. 10.1111/j.1460-9568.1994.tb01008.x.

46. Bussey, T.J., and Saksida, L.M. (2005). Object memory and perception in the medial temporal lobe: An alternative approach. Current Opinion in Neurobiology 15, 730–737. 10.1016/j.conb.2005.10.014.

47. Bussey, T.J., Saksida, L.M., and Murray, E.A. (2006). Perirhinal cortex and feature-ambiguous discriminations. Learning & Memory 13, 103–105. 10.1101/lm.163606.

48. Bartko, S.J., Winters, B.D., Cowell, R.A., Saksida, L.M., and Bussey, T.J. (2007). Perceptual Functions of Perirhinal Cortex in Rats: Zero-Delay Object Recognition and Simultaneous Oddity Discriminations. Journal of Neuroscience 27, 2548–2559. 10.1523/JNEUROSCI.5171-06.2007.

49. Ahn, J.-R., and Lee, I. (2017). Neural Correlates of Both Perception and Memory for Objects in the Rodent Perirhinal Cortex. Cerebral Cortex, 1–13. 10.1093/cercor/bhx093.

50. Hwang, J., and Romanski, L.M. (2015). Prefrontal neuronal responses during audiovisual mnemonic processing. Journal of Neuroscience 35, 960–971. 10.1523/JNEUROSCI.1328-14.2015.

51. Peng, X., and Burwell, R.D. (2021). Beyond the hippocampus: The role of parahippocampal-prefrontal communication in context-modulated behavior. Neurobiology of Learning and Memory 185, 107520. 10.1016/j.nlm.2021.107520.

52. Van Groen, T., and Wyss, J.M. (1990). Extrinsic projections from area CA1 of the rat hippocampus: Olfactory, cortical, subcortical, and bilateral hippocampal formation projections. Journal of Comparative Neurology 302, 515–528. 10.1002/cne.903020308.

53. Quiroga, R.Q., Reddy, L., Kreiman, G., Koch, C., and Fried, I. (2005). Invariant visual representation by single neurons in the human brain. Nature 435, 1102–1107. 10.1038/nature03687.

54. Quiroga, R.Q., Kraskov, A., Koch, C., and Fried, I. (2009). Explicit Encoding of Multimodal Percepts by Single Neurons in the Human Brain. Current Biology 19, 1308–1313. 10.1016/j.cub.2009.06.060.

55. Quiroga, R.Q. (2012). Concept cells: the building blocks of declarative memory functions. Nature Neuroscience 13, 587–597. 10.1038/nrn3251.

56. Reber, T.P., Bausch, M., Mackay, S., Boström, J., Elger, C.E., and Mormann, F. (2019). Representation of abstract semantic knowledge in populations of human single neurons in the medial temporal lobe. PLoS Biol 17, e3000290. 10.1371/journal.pbio.3000290.

57. Tyree, T.J., Metke, M., and Miller, C.T. (2023). Cross-modal representation of identity in the primate hippocampus. Science 382, 417–423. 10.1126/science.adf0460.

58. Yassa, M.A., and Stark, C.E.L. (2011). Pattern separation in the hippocampus. Trends in Neurosciences 34, 515–525. 10.1016/j.tins.2011.06.006.

59. Kent, B.A., Hvoslef-Eide, M., Saksida, L.M., and Bussey, T.J. (2016). The representational-hierarchical view of pattern separation: Not just hippocampus, not just space, not just memory? Neurobiology of Learning and Memory 129, 99–106. 10.1016/j.nlm.2016.01.006.

60. Marr, D., and Brindley, G.S. (1971). Simple memory: a theory for archicortex. Philosophical Transactions of the Royal Society of London. B, Biological Sciences 262, 23–81. 10.1098/rstb.1971.0078.

61. Leutgeb, J.K., Leutgeb, S., Moser, M.-B., and Moser, E.I. (2007). Pattern Separation in the Dentate Gyrus and CA3 of the Hippocampus. Science 315, 961–966. 10.1126/science.1135801.

62. Ahn, J.-R., Lee, H.-W., and Lee, I. (2019). Rhythmic Pruning of Perceptual Noise for Object Representation in the Hippocampus and Perirhinal Cortex in Rats. Cell Reports 26, 2362–2376.e4. 10.1016/j.celrep.2019.02.010.

63. Kaplan, J.T., Man, K., and Greening, S.G. (2015). Multivariate cross-classification: applying machine learning techniques to characterize abstraction in neural representations. Frontiers in Human Neuroscience 9, 1–12. 10.3389/fnhum.2015.00151.

