## Supplemental information for "Neural correlates of modality-specific and modality-invariant object recognition in the perirhinal cortex"

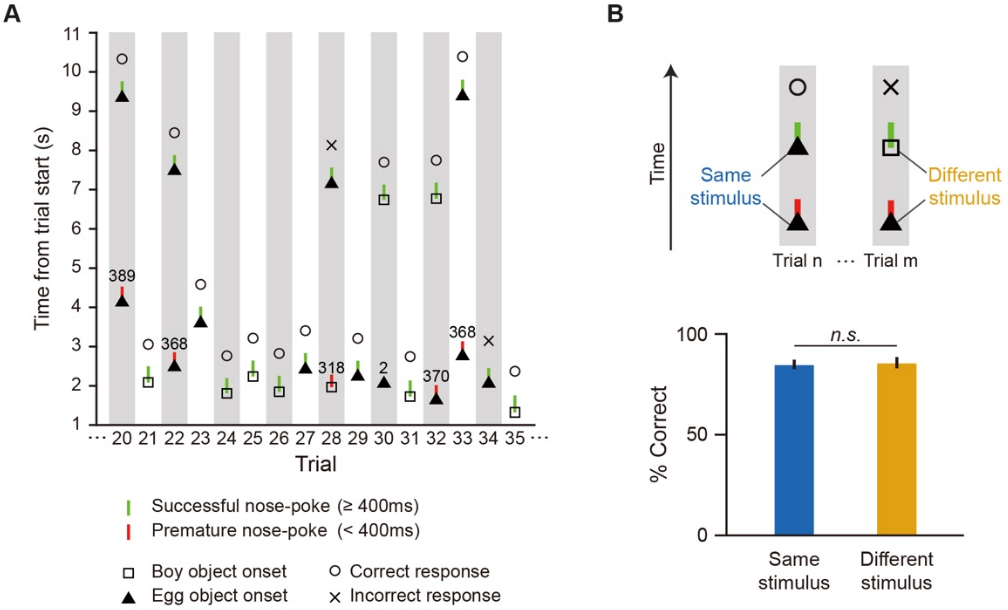


**Fig. S1. Nose-poking behaviors and their influence on performance; related to Fig. 1.** (**A**) Illustration of the task sequence for trials 20 through 35 in an example session. The *Boy* (square) or *Egg* (triangle) object was pseudo-randomly presented regardless of whether the nose-poke was successful (>400 ms) on the previous attempt. Successful nose-poke attempts are marked with green lines. Prematurely withdrawn nose-pokes (<400 ms) are marked with red lines; the numbers above them indicate the duration of the rat's nose-pokes in milliseconds. (**B**) Nose-poke failures did not have a significant influence on correctness. After rats failed to maintain nose-poke for 400 ms (red lines), they either experienced the same (blue) or a different (yellow) object on their next nose-poke attempt (top). There was no significant difference in correctness between the same and different stimulus situations ($t_{(5)} = 0.27$, $p = 0.8$, paired t-test). Data are from sessions where rats (n = 6) performed above the learning criterion (correctness > 75%). n.s., not significant.

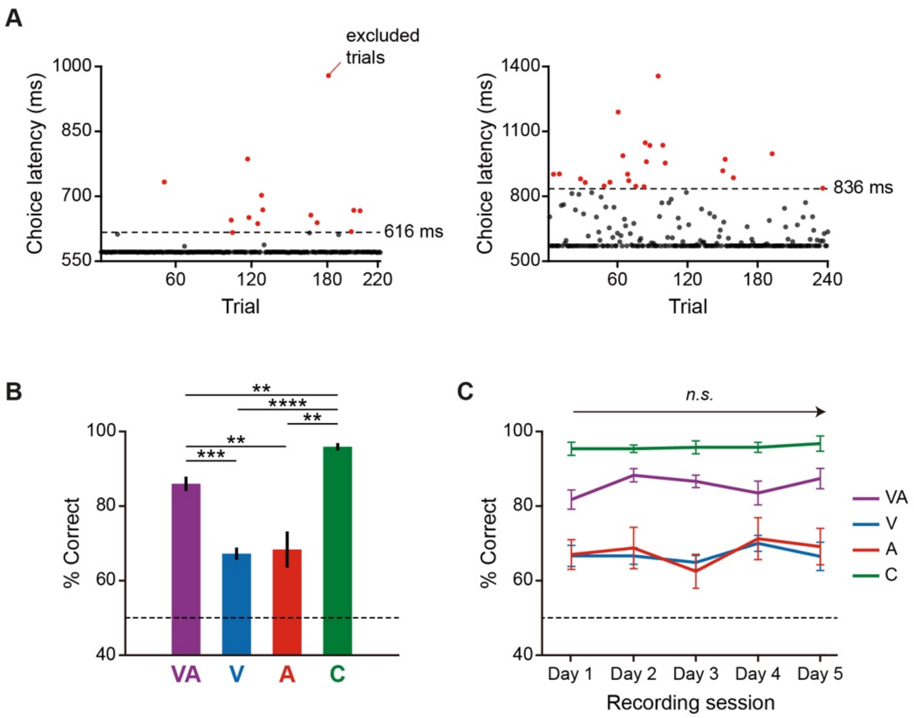


**Fig. S2. Detailed behavioral data from electrophysiological experiments; related to Fig. 3.** (**A**) Choice latency (from cue offset to the end of choice response) data from two example sessions are shown, from rat #10 (left) and rat #13 (right). Each dot indicates the choice latency of each trial. Although choice latency was more stable in rat #10 compared to rat #13, both rats completed most trials within a 600-ms choice latency. Trials were excluded for neural data analysis (red dots) if their choice latency was longer than the median + 3 × the median absolute deviation (dotted lines). (**B**) Average behavioral performance in all recording sessions. There were significant differences in correctness between conditions ($F_{(3,21)}$ = 28.11, $p < 0.0001$; one-way repeated measures ANOVA). Performance in the multimodal condition (VA) was significantly higher than that in visual (V, $t_{(7)} = 7.85$, $p = 0.0005$) and auditory (A, $t_{(7)} = 4.93$, $p = 0.0051$; paired t-test with Holm-Bonferroni correction) conditions. Performance in the control condition (C) was significantly higher than that in all the other conditions (control vs. multimodal, $t_{(7)} = 4.22$, $p = 0.0078$; control vs. visual, $t_{(7)} = 12.51$, $p < 0.0001$; control vs. auditory, $t_{(7)} = 5.93$, $p = 0.0023$; paired t-test with Holm-Bonferroni correction). (**C**) Behavioral performance across recording sessions. Correctness was not significantly different across recording sessions ($F_{(4,28)} = 1.58$, $p = 0.21$; two-way repeated measures ANOVA), indicating that there was no additional learning after repeated sessions. There were significant differences between the conditions ($F_{(3,21)} = 30.04$, $p < 0.0001$), but the interaction effect between the recording session and condition factors was not significant ($F_{(12,84)} = 1.56$, $p = 0.12$; two-way repeated measures ANOVA). Dotted lines indicate chance level performance (50%). Data are presented as means ± SEM ($n = 8$; **$p < 0.01$, ***$p < 0.001$, ****$p < 0.0001$; n.s., not significant).

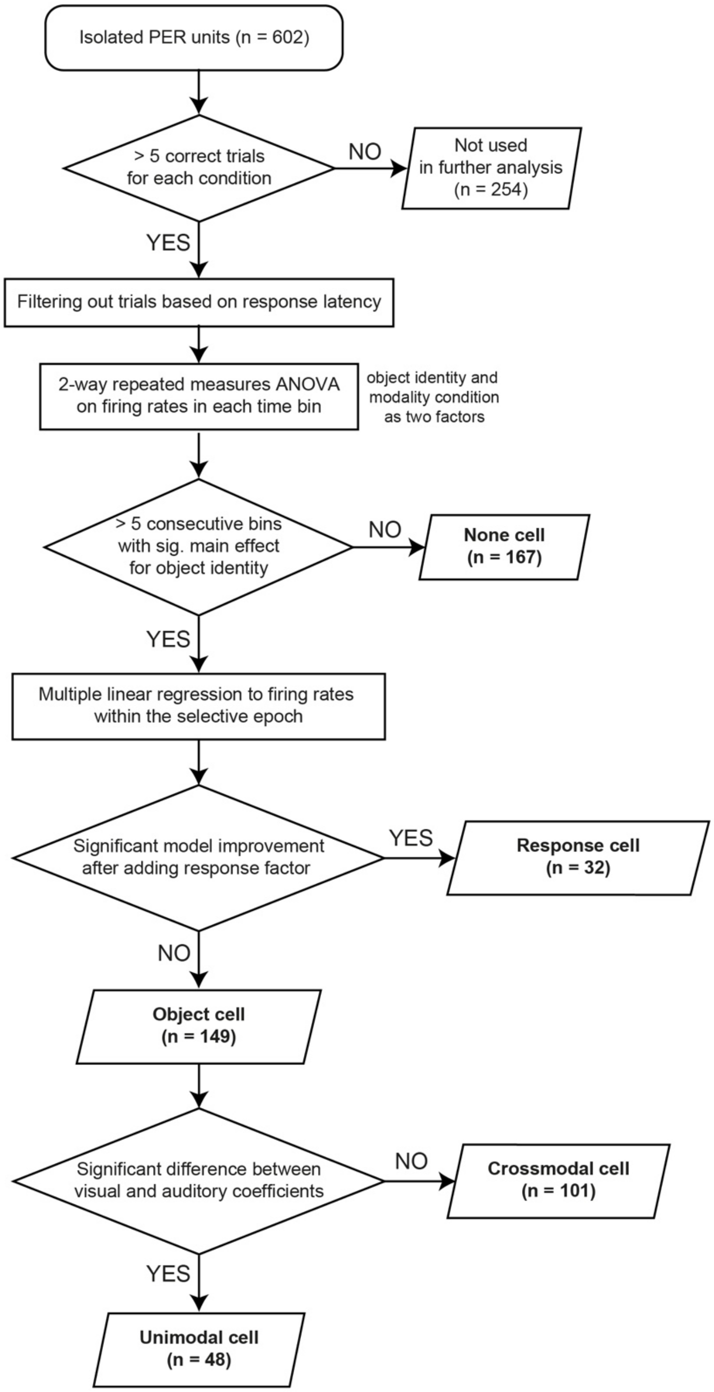


**Fig. S3. Analytic scheme for single-unit data.**

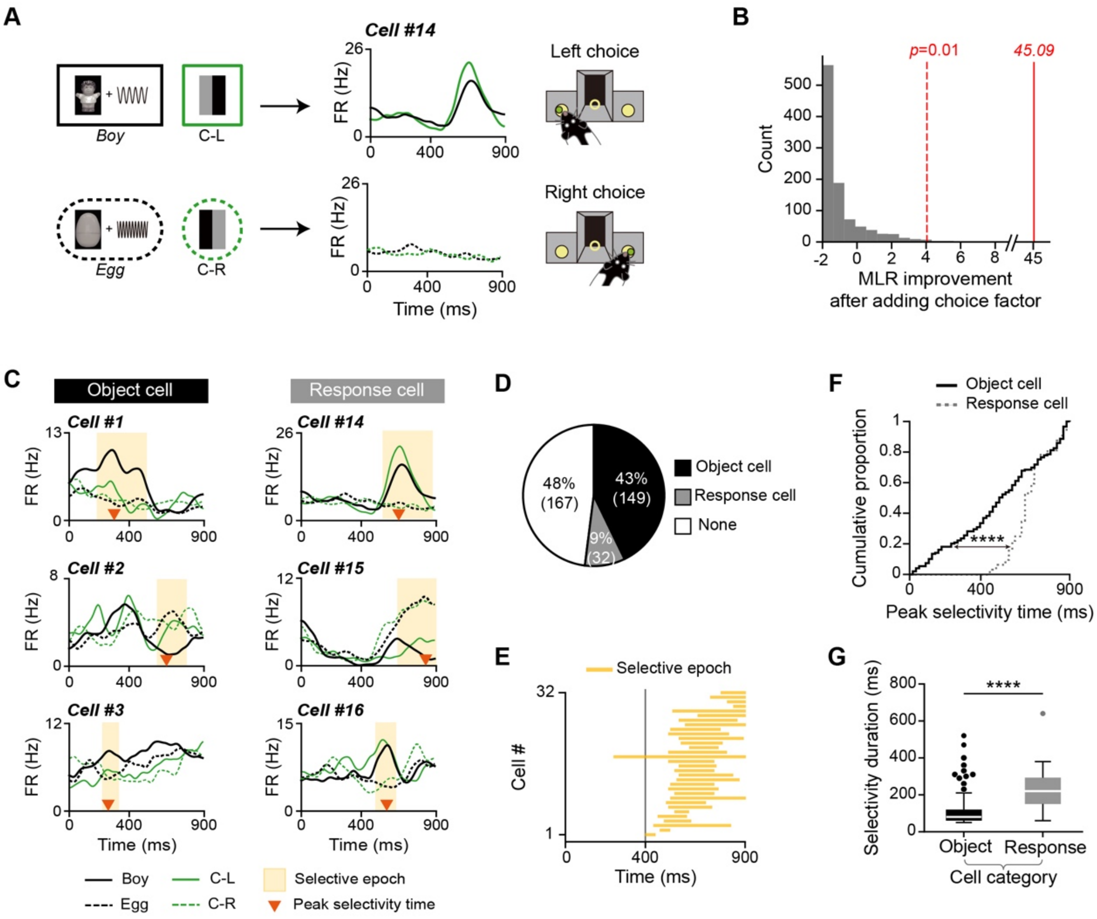


**Fig. S4. Response-selective firing patterns in the PER; related to Fig. 4.** (**A**) Example neuron showing selective firing patterns to a left choice response. Note that object *Boy* and C-L (control-left) conditions were associated with the same left choice response, whereas object *Egg* and C-R (control-right) conditions required the same right choice response. Spike density functions on the left showed increased firing rates when rats were producing a left choice response. (**B**) Classification of the response cell in **A**. The AIC (Akaike Information Criterion) difference was calculated before and after adding the choice factor to the multiple linear regression model (see Methods). The histogram shows the AIC difference calculated from shuffled data (iterations = 1000). Because the cell's actual AIC difference (red solid line) was significantly higher than the alpha level (red dotted line, $p = 0.01$), the neuron was classified as a response cell. (**C**) Examples of object cells (left) and response cells (right) and their firing patterns to object and control conditions. Note that response cells showed overlapping firing patterns to both object and control conditions requiring the same choice response, but object cells did not. (**D**) Proportions of object and response cells within the PER. Numbers in parentheses denote the number of neurons. (**E**)

Population selectivity plot for all response cells in the PER. Most of their selective epochs occurred during the response phase, in contrast to the sequential tiling of the entire task epoch by object cells shown in **Figure 4C**. Gray vertical line indicates the onset of the response phase. (**F**) Cumulative distributions of peak selectivity for object (solid black line) and response (dotted gray line) cell categories. There were significant differences in peak selectivity time between categories (D = 0.46, $p < 0.0001$; Kolmogorov–Smirnov test). The peak selectivity time for most response cells occurred after 400 ms (i.e., the response phase). (**G**) Comparison of the duration of selectivity between object and response cells. The duration of selectivity for the response cell category was significantly longer compared with the object cell category (U = 832, ****$p < 0.0001$; Mann-Whitney U test); n.s., not significant.

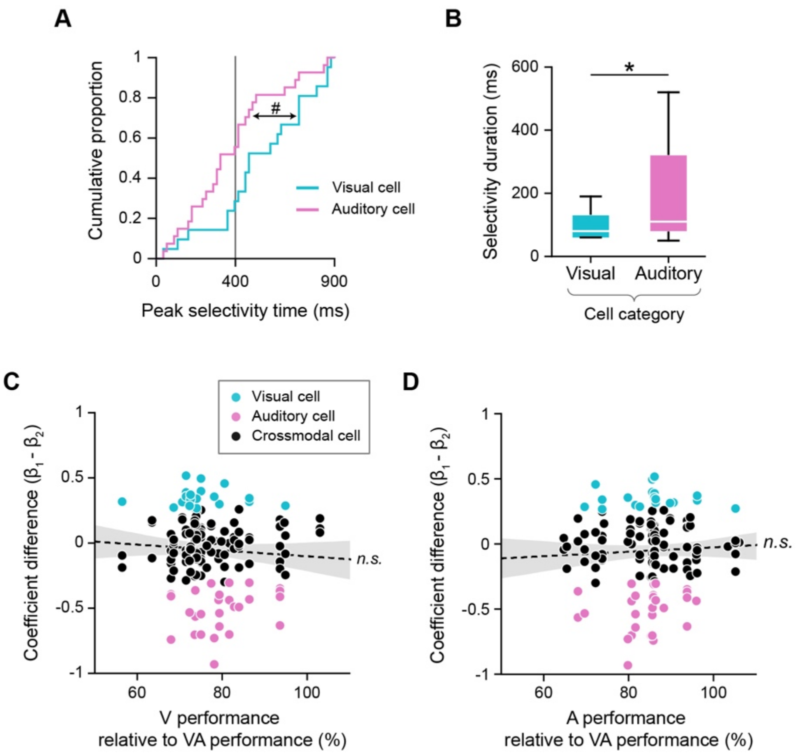


**Fig. S5. Selective firing patterns of unimodal cells and their behavioral correlates.** (**A**) Cumulative distributions of peak selectivity time for visual and auditory cells. Differences in peak selectivity time were marginally significant between the two categories of cells (D = 0.38, $p = 0.053$; Kolmogorov–Smirnov test). Peak selectivity for auditory cells was more likely to occur during the sample phase. The vertical gray line indicates the boundary between sample and response phases. (**B**) Comparison of the duration of selective epochs between visual and auditory cells. The duration of selectivity for auditory cells was significantly longer than that for visual cells (U = 388.5, $p = 0.03$; Mann-Whitney U test). (**C**) Scatter plot of differences in coefficients ($\beta_1 - \beta_2$) showing relative performance in the visual condition, used to investigate the relationship between a neuron's visual preference and the performance in the visual condition. Relative performance was obtained by dividing correctness in the visual (V) condition by correctness in the multimodal (VA) condition. Visual cells (cyan) were present regardless of the rat's performance in the visual condition. There was no significant linear relationship between performance and the difference in coefficients (r = -0.071, $p = 0.39$). (**D**) Relationship between a neuron's auditory preference and performance in the auditory condition. Relative performance was calculated by dividing correctness in the auditory (A) condition by correctness in the multimodal (VA) condition. Auditory cells (pink) were present irrespective of the rat's performance in the auditory condition. No significant relationship was found between the difference in coefficients and relative performance in the auditory condition (r = 0.054, $p = 0.51$). Dotted black lines indicate the linear regression line, and the shaded areas represent the 95% confidence interval. #$p = 0.053$. *$p < 0.05$. n.s., not significant.

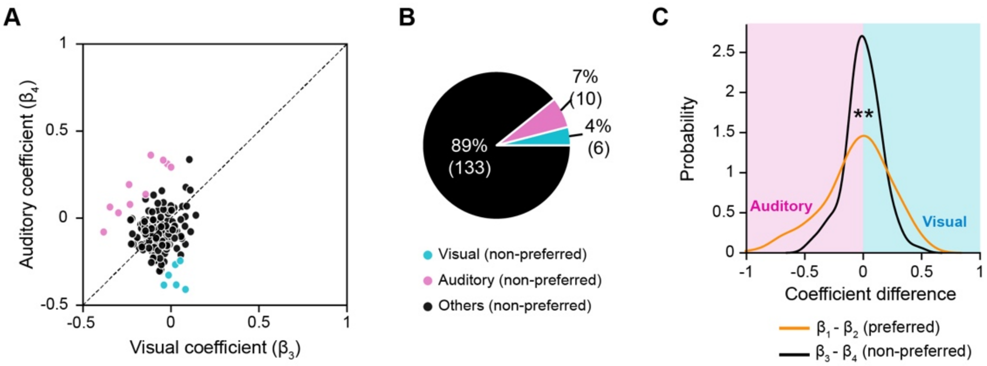


**Fig. S6. Regression coefficients for non-preferred objects; related to Fig. 5.** (**A**) Scatter plot showing the regression coefficients $\beta_3$ and $\beta_4$ for a non-preferred object (i.e., object conditions with lower firing rates). Cells were classified into visual (cyan) or auditory (pink) categories using the same procedure as in **Figure 5**, but with $\beta_3$ and $\beta_4$ instead. $\beta_3$ and $\beta_4$ values for most neurons were around zero, indicating that they were not modulated by the modality information of non-preferred objects. (**B**) Proportions of visual and auditory cells classified using regression coefficients for the non-preferred object. Only a handful of neurons were classified as having a significant preference for the visual or auditory information of the non-preferred object. Numbers in parentheses indicate the number of cells. (**C**) Kernel density estimations of differences in coefficients for preferred (orange) and non-preferred (black) object conditions. For the preferred object, there were more neurons with extremely negative (auditory) or positive (visual) differences in coefficient values. However, the difference in coefficients for the non-preferred object was centered around zero, indicating no modulation by a specific sensory modality. The distributions of coefficient differences were significantly different between preferred and non-preferred object conditions (D = 0.19, $p = 0.007$; Kolmogorov–Smirnov test). $^{**}p < 0.01$.

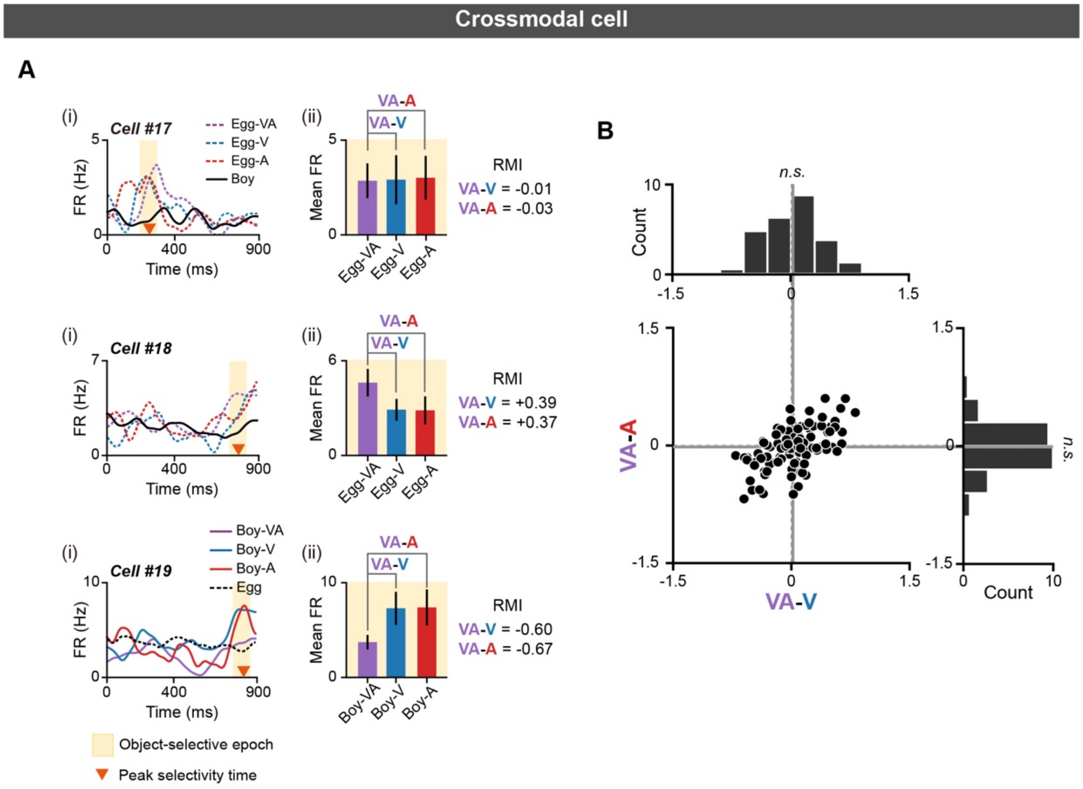


**Fig. S7. Firing rate modulations of crossmodal cells to different modality conditions; related to Fig. 6.** (**A**) Examples of crossmodal cells and their modulation patterns in different modality conditions. Spike density functions (i) and mean firing rates within the selective epoch (ii) demonstrate heterogeneous modulation patterns in crossmodal cells. In cell #17, mean firing rates were similar across all modality conditions of the preferred object (*Egg*), and RMI values were near zero (VA – V = -0.01, VA – A = 0.03). The firing rates of cell #18 were higher in the multimodal condition than in visual or auditory conditions, resulting in positive RMI values (VA – V = 0.39, VA – A = 0.37). On the other hand, firing rates for cell #19 were lower in the multimodal condition compared with both visual and auditory conditions, resulting in negative RMI values (VA – V = -0.6, VA – A = -0.67). (**B**) Scatter plot and histograms of VA – V and VA – A in crossmodal cells. Average VA – V (vertical gray line) and average VA – A (horizontal gray line) were not significantly different from zero (VA – V, $t_{(100)} = 0.7$, $p = 0.49$; VA – A, $t_{(100)} = 0.49$, $p = 0.62$; one-sample t-test); n.s., not significant.

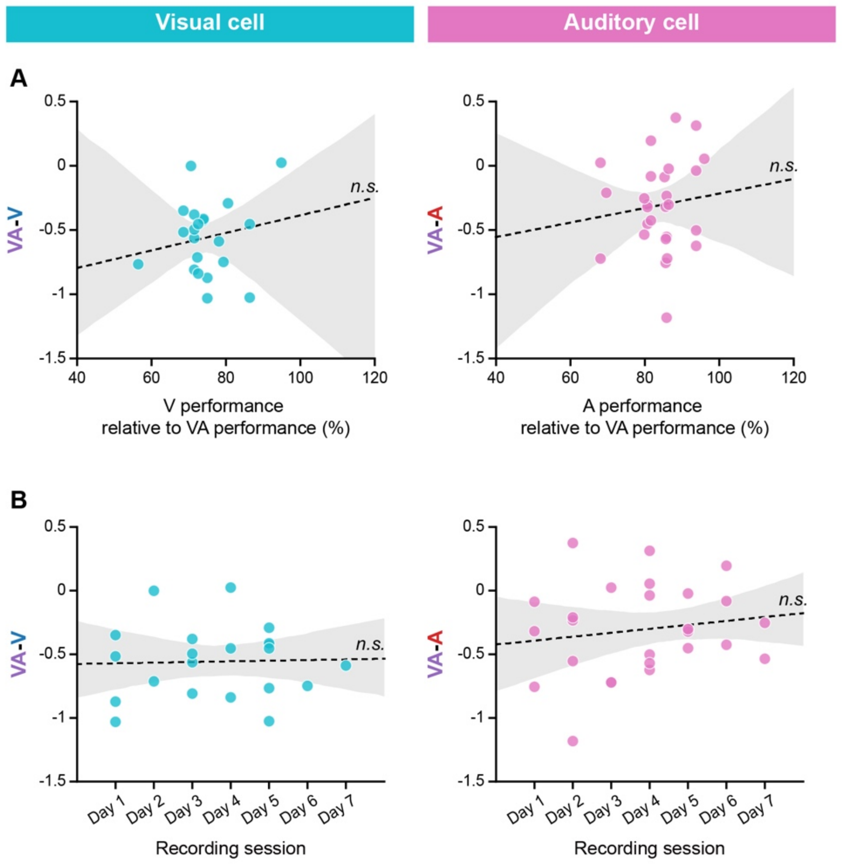


**Fig. S8. Relationship between multisensory suppression and behavior; related to Fig. 6.** (**A**) Scatter plots of RMI values (VA – V or VA – A) and relative performance, displayed separately for visual (left) and auditory (right) cells, used to determine whether multisensory suppression (i.e., negative VA – V or VA – A values) in visual or auditory cells is related to lower correctness in visual or auditory conditions. Relative performance was obtained by dividing correctness in the visual (V) or auditory (A) condition by correctness in the multimodal (VA) condition. Neither VA – V nor VA – A became more negative as the rat performance worsened in the visual or auditory condition. In both cell categories, no significant linear relationship was found between relative performance and RMI values (visual, r = 0.19, $p = 0.42$; auditory, r = 0.12, $p = 0.57$). Each dot indicates individual visual or auditory cells. (**B**) Scatter plots of RMI values (VA – V or VA – A) and neurons' recorded sessions, displayed separately for visual (left) and auditory (right) cells, used to investigate whether multisensory suppression (i.e., negative VA – V or VA – A values) in visual or auditory cells is related to the novelty of the visual or auditory condition. Even on days 5 through 7, when rats were sufficiently acclimated to visual or auditory conditions, neurons exhibited negative VA – V or VA – A values. Therefore, it is unlikely that the suppression of activities in the multimodal condition is attributable to repetition suppression in the familiar multimodal condition. In both cell categories, no significant linear relationship was found between the recording session and RMI (visual, r = 0.03, $p = 0.89$; auditory, r = 0.16, $p = 0.44$). The dotted black lines indicate the linear regression line, and the shaded areas represent the 95% confidence interval. n.s., not significant.

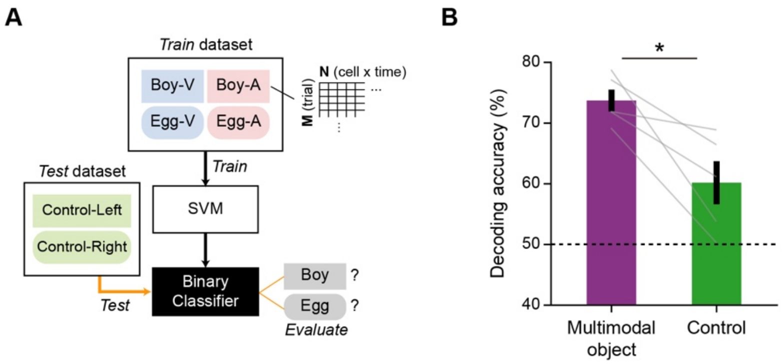


**Fig. S9. Control analysis of modality-invariant object decoding; related to Fig. 7.** (**A**) If modality-invariant decoding was successful simply because an object always required the same choice response, we would expect to observe comparable decoding accuracy for control stimuli requiring the same choice response. We therefore performed decoding of control stimuli based on visual and auditory object conditions using a linear support vector machine (SVM). The classifier was trained using the same dataset as used for modality-invariant decoding in **Figure 7D**. However, this time, we tested whether the same classifier could discriminate between two control stimuli that required the same choice response instead of the multimodal objects. (**B**) Comparison of decoding accuracies for multimodal objects (**Fig. 7D**) and control stimuli. The decoding accuracy of the control stimuli was not comparable to that of the multimodal objects, suggesting that modality-invariant decoding was enabled by the object identity rather than the choice response. Decoding accuracy was significantly different between the two decoding methods ($t_{(4)} = 3.61$, $p = 0.023$, paired t-test). Dotted black lines indicate the chance level of decoding accuracy obtained from surrogate data. Data are presented as means ± SEM (*$p < 0.05$).
